# Protein-intrinsic properties and context-dependent effects regulate pioneer-factor binding and function

**DOI:** 10.1101/2023.03.18.533281

**Authors:** Tyler J. Gibson, Melissa M. Harrison

## Abstract

Chromatin is a barrier to the binding of many transcription factors. By contrast, pioneer factors access nucleosomal targets and promote chromatin opening. Despite binding to target motifs in closed chromatin, many pioneer factors display cell-type specific binding and activity. The mechanisms governing pioneer-factor occupancy and the relationship between chromatin occupancy and opening remain unclear. We studied three *Drosophila* transcription factors with distinct DNA-binding domains and biological functions: Zelda, Grainy head, and Twist. We demonstrated that the level of chromatin occupancy is a key determinant of pioneering activity. Multiple factors regulate occupancy, including motif content, local chromatin, and protein concentration. Regions outside the DNA-binding domain are required for binding and chromatin opening. Our results show that pioneering activity is not a binary feature intrinsic to a protein but occurs on a spectrum and is regulated by a variety of protein-intrinsic and cell-type-specific features.

## Introduction

During animal development, the totipotent embryo divides and differentiates to give rise to the diverse cell and tissue types of the adult organism. The establishment of cell-type specific gene expression programs is driven by transcription factors (TFs) that bind sequence specifically to *cis*-regulatory elements. The packaging of DNA into chromatin obstructs TFs from accessing their target motifs^1^. Thus, many TFs only bind the small subset of genomic motifs that fall within regions of open chromatin^2^. A specialized subset of transcription factors, termed pioneer factors (PFs), are defined by their ability to bind target motifs on nucleosomes and promote chromatin opening, allowing additional TFs to bind these newly accessible motifs and activate transcription^3,4^. Through this activity, PFs drive cell-type specific gene expression programs that determine cell fate. PFs are essential players in development, induced reprogramming, and disease^5^.

Despite their ability to target closed, nucleosomally occupied binding sites, PFs do not bind to all genomic instances of their target motifs. PFs also have cell-type specific patterns of binding and activity, suggesting that features beyond sequence motif govern PF function^6–11^. Chromatin structure may influence PF binding as cell-type specific occupancy is often associated with distinct chromatin features^10–12^. Many PFs bind primarily to naïve chromatin that is inactive and lacks most histone modifications^6,7^. Together, these studies indicate that PF occupancy is regulated by both genomic context and cell-type specific features. Nonetheless, the mechanisms that govern PF occupancy and activity remain largely unknown.

PF activity is regulated at a step other than chromatin binding. PFs are defined by their ability to promote chromatin opening, yet many PFs are only required for chromatin accessibility at a subset of their binding sites. For example, upon depletion of the PFs Zelda, GAGA-factor, Oct4, or Sox2, chromatin accessibility is lost only at a subset of the binding sites for the respective factor^13–15^. Pax7, a PF important for specification of pituitary melanotropes, binds to thousands of “pioneer primed” regions where binding does not result in chromatin opening^12^. Further separating PF binding from opening, at some Pax7-bound loci there is a temporal delay between binding and chromatin opening. This distinction between binding and opening may be due to PFs recruiting additional factors to drive chromatin accessibility. For example, both GAGA factor and Oct4 require nucleosome remodelers to promote chromatin accessibility^16,17^. Nonetheless, PF binding can also perturb nucleosome structure. Oct4 binding to nucleosomes results in local remodeling of histone-DNA interactions, especially at the nucleosome entry-exit site^18^. Together, these observations highlight that the relationship between PF binding and activity is not well understood.

To begin to elucidate the factors that regulate PF binding and activity, we focused on two well-studied PFs that control conserved developmental transitions, Zelda (Zld) and Grainy head (Grh). Zld is essential for driving early embryonic development in *Drosophila*^19,20^. Immediately following fertilization, the zygotic genome is transcriptionally silent, and development is controlled by maternally deposited mRNAs and proteins. Maternal products are gradually degraded as the zygotic genome is activated during a process known as the maternal-to-zygotic transition (MZT)^21,22^. In *Drosophila*, the pioneering activity of Zld is essential for activating transcription from the zygotic genome^13,19,23,24^. Since the discovery of Zld as an activator of the initial wave of zygotic transcription, pioneer factors have been shown to drive zygotic genome activation in all other species studied to date^25–29^. Similar to the role of Zld in genomic reprogramming in the early embryo, Zld promotes the neural stem cell fate in the developing larval brain^30^. Despite the shared ability to promote the undifferentiated fate in both tissues, the majority of Zld-binding sites are unique to either tissue. In contrast to the tissue-specific chromatin occupancy of Zld, Grh-binding sites are largely shared between tissues^31^. Grh is a master regulator of epithelial cell fate that is conserved among metazoans and functions as a pioneer factor in both *Drosophila* larva and mammalian cell culture^32–35^. However, in the early *Drosophila* embryo Grh is dispensible for chromatin accessibility, and only displays pioneering activity in late embryos or larval tissues. Thus, Grh pioneering activity, but not Grh binding is regulated over development.

The essential roles of Zld and Grh in diverse biological processes and the context-dependent binding and activity of these factors makes them excellent models to explore how PF binding is regulated and how PF occupancy relates to chromatin opening. Because both Zld and Grh are expressed immediately following fertilization, previous studies have largely relied on loss-of-function experiments. Thus, it has been difficult to disentangle causal relationships between chromatin structure and PF activity. We therefore leveraged the well-characterized Drosophila Schneider 2 (S2) cell-culture system in which neither Zld nor Grh is endogenously expressed as a platform to identify features that promote or prevent PF binding and activity. When ectopically expressed, Zld and Grh bound to closed chromatin, established chromatin accessibility, and promoted transcriptional activation at a subset of binding sites. By comparing the features of these factors and relating exogenous activity to *in vivo* functionality, we demonstrated that pioneering depended on a combination of local chromatin structure, sequence context, and PF concentration. We propose that pioneering activity is not a binary feature intrinsic to a protein, but occurs on a spectrum and can be regulated by a variety of protein-intrinsic and cell-type specific features.

## Results

### Ectopically expressed pioneer factors bind and open closed chromatin

To investigate the mechanisms by which pioneer factors bind and open chromatin, we generated stable S2 cell lines capable of individually expressing either Zld or Grh (Extended Data Fig. 1a). These cell lines allowed the inducible and tunable expression of each factor, as neither pioneer factor is normally expressed in wild-type S2 cells (Extended Data Fig. 1b-d). We expressed each factor at approximately physiological levels and determined genome-wide binding sites using chromatin immunoprecipitation coupled with sequencing (ChIP-seq). We identified 9203 peaks for Zld and 13851 for Grh. Additional ChIP experiments were performed in wild-type cells using anti-Zld, or anti-Grh to control for antibody specificity. Despite undetectable levels of protein by immunoblot, ChIP signal for both proteins was identified in wild-type cells (Extended Data Fig 1e-f). To ensure that all peaks used for subsequent analysis reflected those that were gained upon induction, we used these wild-type ChIP datasets as controls for peak calling.

Having established that Zld and Grh bind thousands of sites in S2 cells, we determined chromatin accessibility in wild-type cells or those expressing Zld or Grh using assay for transposase-accessible chromatin (ATAC)-seq. Comparisons between Zld- or Grh-expressing cells and wild-type cells identified 6546 differentially accessible regions for Zld (3769 increased, 2777 decreased) and 13732 for Grh (4784 increased, 8948 decreased) (Extended Data Fig. 2a,d)^36^. We integrated our ChIP-seq and ATAC-seq data to determine the connection between binding and accessibility. We identified three classes of PF-bound sites (Fig. 1a,b; Supplementary Table 1). Class I sites are accessible in wild-type cells and remain accessible when bound by the ectopically expressed factor. Class II sites are inaccessible in wild-type cells. They are bound by Zld or Grh but this binding does not lead to chromatin opening. Class III sites are inaccessible in wild-type cells, but become accessible upon PF binding. The majority of ChIP-seq peaks for Zld and Grh were class I sites. For each factor, only a subset of binding sites were within closed chromatin, and at only a subset of these did the chromatin become accessible (Fig. 1c,f). Class I binding sites occurred predominantly at promoters, while the majority of class II and III sites were at promoter-distal sites (Fig. 1d,g). Genome-wide analysis of signal intensities at all class II and III sites confirmed that these sites lack ATAC signal in wild-type cells and that class III sites undergo robust chromatin opening upon Zld or Grh induction (Fig. 1e,h). Together, these data show that when expressed exogenously at physiolgical levels, Zld and Grh bind and open closed chromatin at a subset of binding sites.

**Figure 1:**
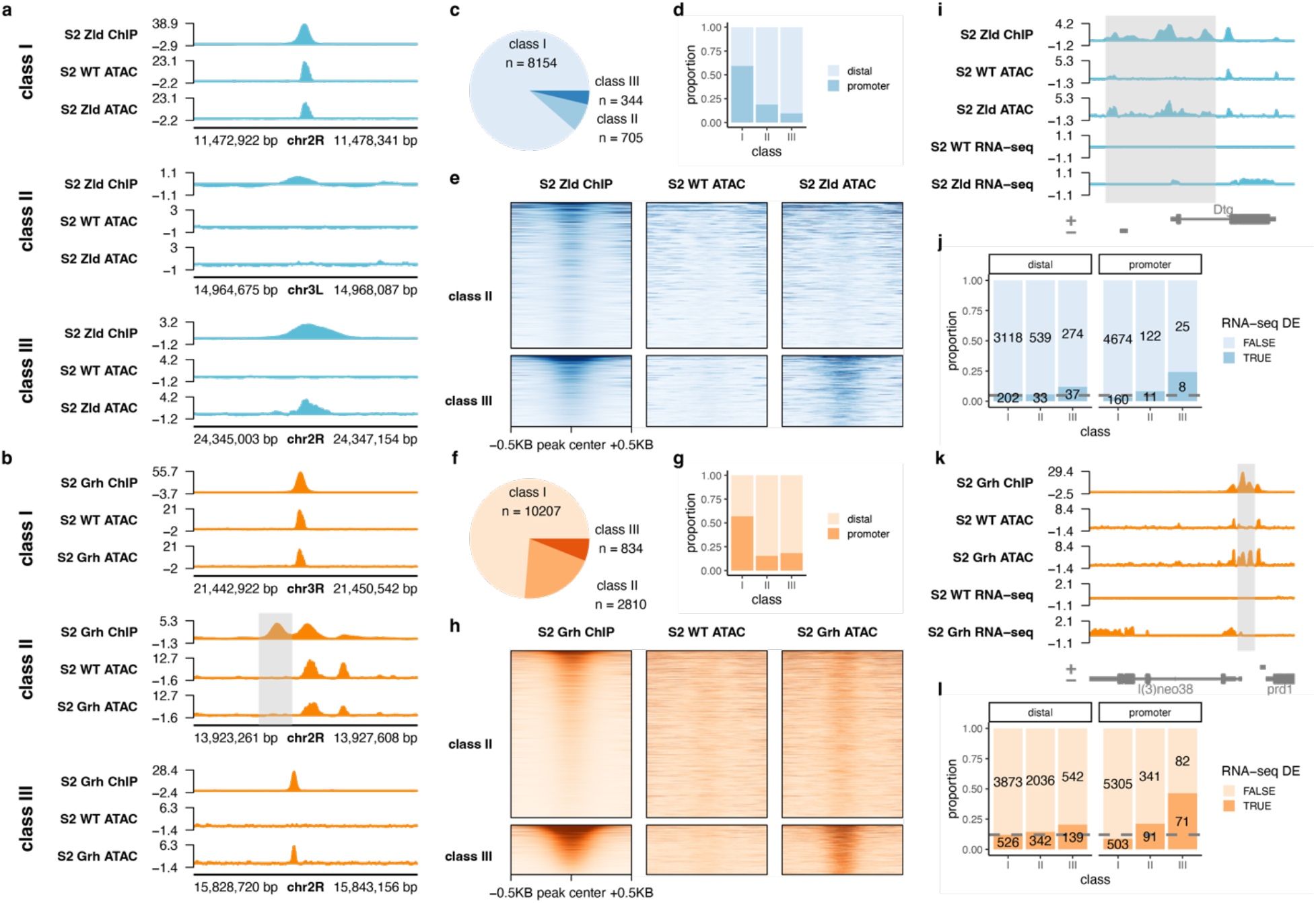
Ectopically expressed pioneer factors open chromatin and activate transcription. **a,b,** Genome browser tracks showing examples of individual class I, II or III regions for Zld **(a)** or Grh **(b)**. ChIP seq and ATAC-seq signal are shown for Zld- or Grh-expressing cells along with ATAC-seq signal for wild-type (WT) cells lacking Zld or Grh expression. **c,f,** Pie charts showing the distribution of class I, II, and III binding sites for Zld **(c)** or Grh **(f)**. **d,g,** The proportion of class I, II or III binding sites that occur at promoters (-500 to +100 bp around transcription start site) or promoter distal regions for Zld **(d)** or Grh **(g)**. **e,h,** Heatmaps showing ChIP-seq and ATAC-seq signal at class II and III regions for Zld **(e)** or Grh **(h). i,k,** Example genome browser tracks showing ChIP-seq, ATAC-seq and RNA-seq signal at class III regions (shaded area) where changes in chromatin accessibility (shaded region) are associated with changes in gene expression for Zld **(i)** or Grh **(k)**. **j,l,** Proportion of class I, II or III regions that is proximal to a gene that is differentially expressed (DE) upon Zld **(j)** or Grh **(l)** expression. Promoter-proximal and promoter-distal binding sites are shown separately. The grey dotted line indicates the percentage of all binding sites that are proximal to a differentially expressed gene.

Next, we sought to determine the binding dynamics of Zld and Grh. Zld and Grh protein was detectable at 4 or 12 hours following induction, respectively (Extended Data Fig. 3a-b). We used CUT&RUN to measure Zld and Grh binding to chromatin at 0, 4, 12, 24 or 48 hours following induction^37^. Both proteins were bound to class I regions even in uninduced cells (0H), suggesting that these sites can be occupied even when protein levels are below the limit of detection of immunoblotting (Extended Data Fig 3c-d). Class II and III regions were bound at 4 hours, with slight increases in occupancy at later time points, demonstrating that Zld and Grh bind closed chromatin rapidly following induction of protein expression.

### Chromatin opening correlates with transcriptional activation

To identify whether Zld- and Grh-mediated chromatin accessibility led to transcriptional activation, we performed RNA-seq. Differential expression analysis comparing induced cells to wild-type cells identified 505 differentially expressed genes for Zld and 1073 for Grh (Extended Data Fig. 2b,e). Genes activated by Zld were enriched for the gene ontology (GO) term embryonic morphogenesis, consistent with the known function of Zld^38^, as well as imaginal disc development, in which Zld-target genes have not been well characterized (Extended Data Fig. 2g). The genes activated by Grh were enriched for GO terms related to cuticle and epithelial development, consistent with the endogenous function of Grh (Extended Data Fig. 2h)^31^.

We compared our RNA-seq and ATAC-seq data, allowing us to investigate the relationship between chromatin opening and transcriptional activation. Overall, chromatin opening was correlated with transcriptional activation (Fig. 1i-l, Extended Data Fig. 2c,f). When we individually analyzed the three classes of binding sites for changes in gene expression, we found that class III sites were most likely to be proximal to upregulated genes (Fig. 1j,l). This was particularly evident at class III promoters. Thus, PF-dependent increases in chromatin accessibility were correlated with transcriptional activation, although binding and chromatin opening did not always result in gene activation.

### Twist binds extensively to closed chromatin and opens chromatin at a subset of sites

Zld and Grh had previously been shown to have features of PFs^13,23,24,33,35^. To determine how other factors would behave in our system, we tested an additional transcription factor, Twist (Twi). Twi is a master regulator of mesodermal cell fate^39^ and, importantly, was not expressed in our wild-type S2 cells. Like Zld and Grh, Twi is an important regulator of *Drosophila* embryonic development. However, Twi has properties that are distinct from these PFs. Unlike Grh, many Twi-binding sites are developmentally dynamic and are not maintained through embryonic development^40^. Additionally, Twi binding in the early embryo requires Zld pioneering activity, as mutations to Zld-binding sites disrupted Twi binding to the *cactus* enhancer^41^. Deep learning models trained on ChIP-seq and ATAC-seq data for Zld and Twi suggest that Zld motifs contribute to chromatin accessibility and Twi binding, while Twi motifs are not predictive of chromatin accessibility^42^. To test the properties of Twi in our system, we generated stable cell lines that inducibly express Twi at approximately physiological levels (Extended Data Fig. 4a). As we had done for Zld and Grh, we determined Twi binding and activity using ChIP-seq, ATAC-seq, and RNA-seq. These data revealed the same three classes of binding sites as described for Zld and Grh (Fig 2a,b; Extended Data Fig. 1g). Class I binding sites were enriched at promoters, while class II and III binding sites were mostly promoter-distal (Fig 2c). Twi bound extensively to closed chromatin, with 27.8% of binding sites lacking detectable chromatin accessibility (Fig. 2b). A relatively small subset of these sites were class III, where binding to closed chromatin resulted in chromatin opening (Fig 2a-d; Extended Data Fig. 4b). RNA-seq revealed that Twi activated transcription of hundreds of genes (Extended Data Fig. 4c). GO analysis demonstrated that genes upregulated upon Twi expression are enriched for mesodermal genes, consistent with the endogenous function of Twi (Extended Data Fig. 4d)^40^. Despite Twi possessing features distinct from Zld and Grh *in vivo*, ectopically expressed Twi bound closed chromatin and promoted accessibility.

**Figure 2:**
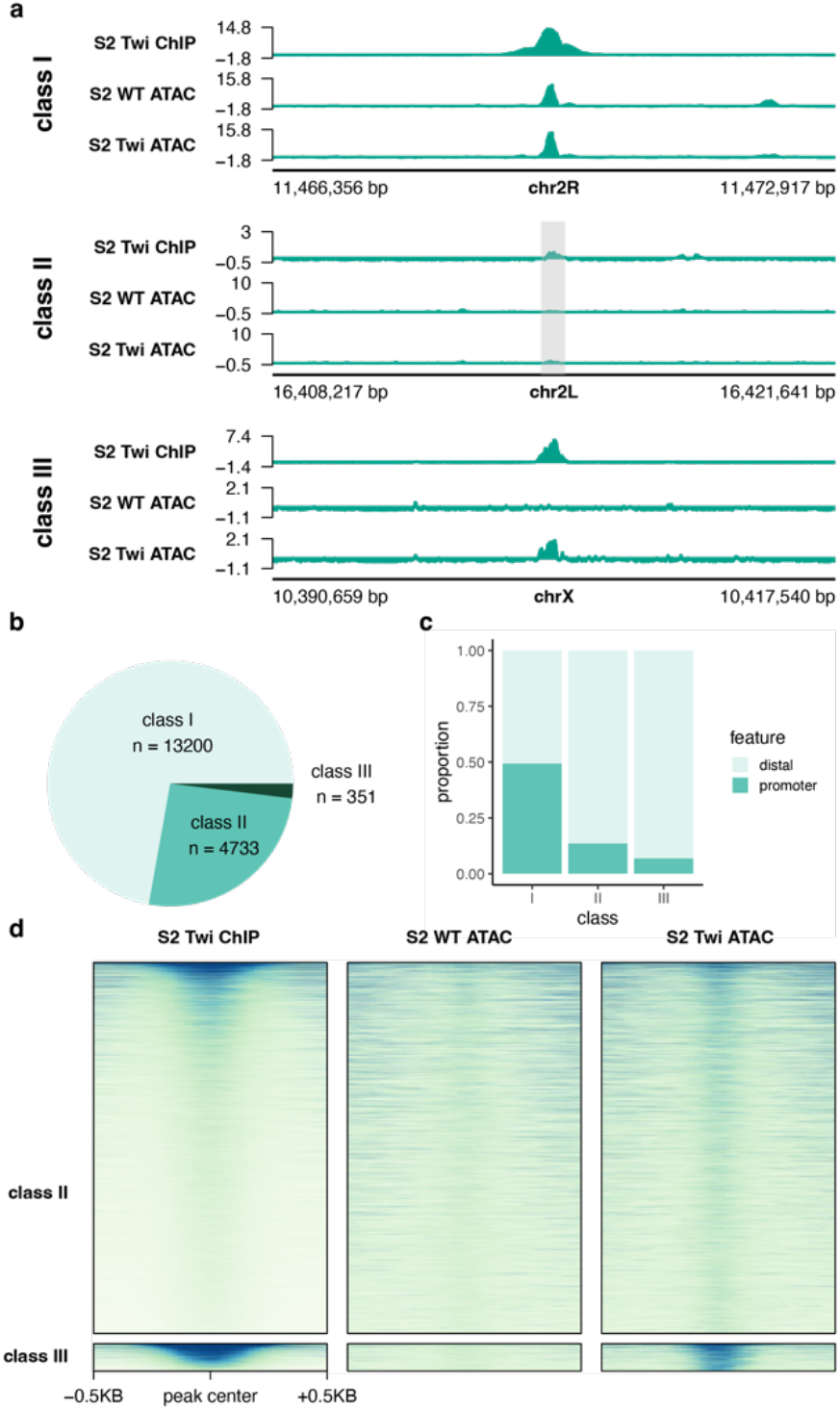
Twist binds closed chromatin extensively and drives accessibility at a limited number of sites. **a,** Genome browser tracks showing examples of individual class I, II or III regions **b,** Pie charts showing the distribution of class I, II, and III binding sites. **c,** The proportion of class I, II or III binding sites that occur at promoters (-500 to +100 bp around transcription start site) or promoter distal regions. **d,** Heatmap showing ChIP-seq and ATAC-seq signal at class II and III regions.

### Ectopically expressed transcription factors bind opportunistically to active chromatin

Comparing binding sites for Zld, Grh, and Twi showed that many class I sites were common to all three factors (Fig. 3a). This contrasted with class II and III peaks, which were largely specific to each individual factor. Control IP experiments in wild-type S2 cells support that these peaks are not due to nonspecific immunoprecipitation (Extended Data Fig. 1e-g). We hypothesized that the overlapping class I binding sites may occur at active *cis*-regulatory regions in wild-type cells. To test this, we analyzed published ChIP-seq data from S2 cells for markers of active chromatin: H3K27ac, CBP, H3K4me1, H3K4me3, and H2AV^43–45^. Overlapping class I regions showed high levels of these chromatin marks (Fig. 3b), suggesting that exogenously expressed factors may bind nonspecifically to active regions of the genome.

**Figure 3:**
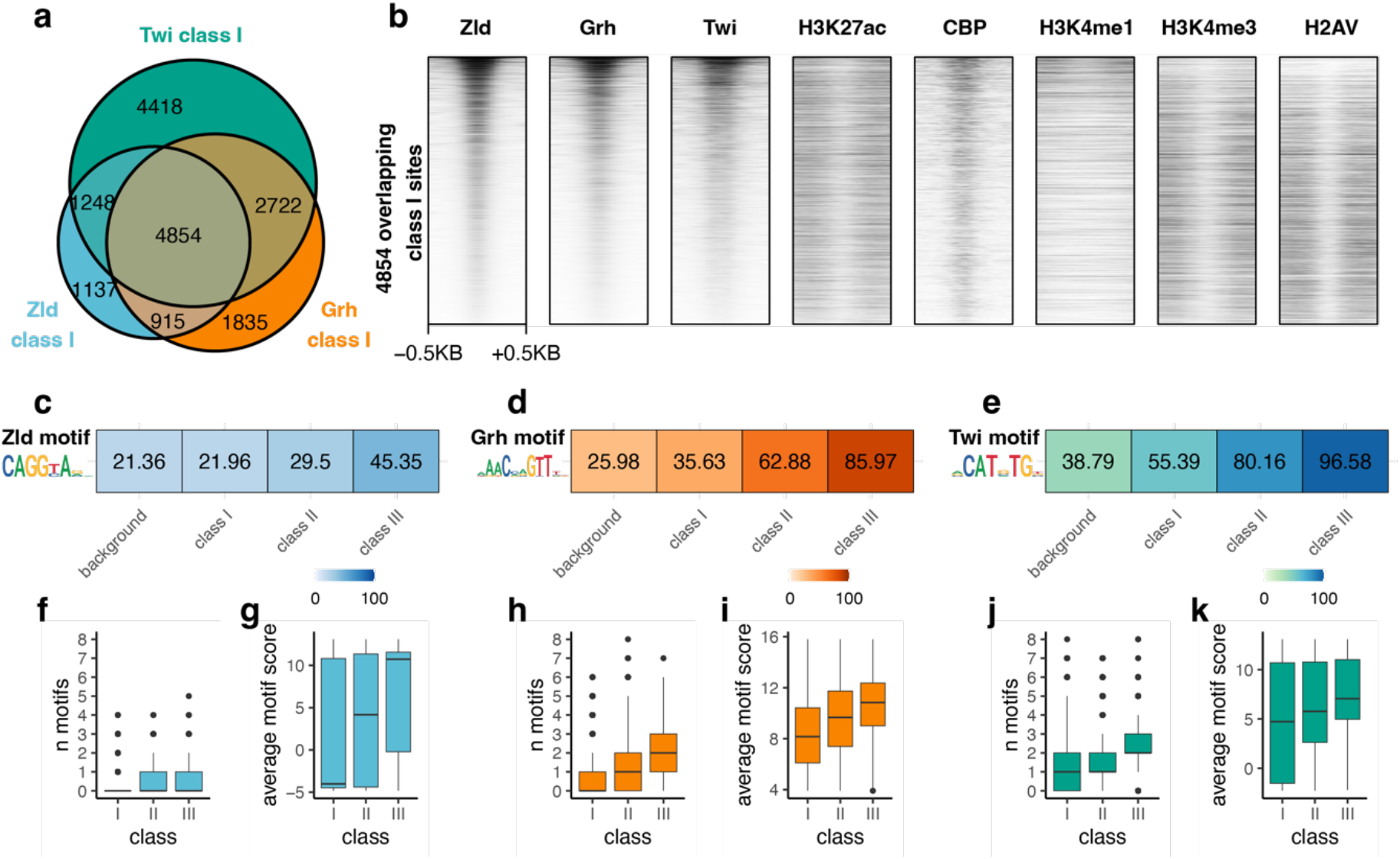
Motif content shapes pioneer-factor activity. **a,** Venn diagram showing overlap between class I regions for Zld, Grh and Twi. **b,** Heatmap of ChIP-seq signal at class I sites that overlap between Zld, Grh, and Twi. Zld, Grh, and Twi ChIP-seq data were generated in this study. Remaining ChIP-seq datasets were previously published (see Supplementary table 2)^43–45^. **c-e,** Heatmaps showing the percentage of regions containing a canonical Zld **(c)**, Grh **(d)**, or Twi motif **(e)**. Logos plot of the canonical motif are shown alongside the heatmap. Percentages are shown for class I, II or III binding sites, or all wild-type ATAC-seq peaks as a control for the background motif frequency within regulatory elements. **f,h,j,** Boxplots of the average number of motifs per peak in class I, II, or III regions for Zld **(f)**, Grh **(h)**, or Twi **(j)**. **g,i,k,** Boxplots of the average motif log-odds score for class I, II or III regions for Zld **(g)**, Grh **(i)**, or Twi **(k)**. Motif scores were determined by comparing each identified motif instance to the canonical motif. For all boxplots, line shows the median, boxes extend from the 25th to the 75th percentile, and whiskers show 1.5 × the interquartile range. Outlier points beyond the range of the whiskers are shown individually.

Considering the enrichment of marks of active chromatin at class I sites, we tested if there were any features of chromatin distinctive to class II or III regions. Analysis of published ChIP-seq data demonstrated that class II and III sites have low levels of many histone modifications, chromatin-associated proteins, and transcription factors (Extended Data Fig. 5a,c,e)^43–55^. This supports the hypothesis that PFs preferentially bind naïve chromatin that is inaccessible and devoid of most histone modifications^6^. The ability of TF binding to promote chromatin accessibility at class III sites and not class II sites was not explained by a pre-existing lack of histones; the levels of H3 and H1 were higher at class III than at class II sites (Extended Data Fig. 5a,c,e). While most class III sites were regions of naïve chromatin, a subset were within regions marked with the repressive histone modification H3K27me3 (Extended Data Fig. 5b,d,f). In fact, H3K27me3 levels were higher in class III compared to class I or II regions (Extended Data Fig. 5a,c,e). Together these analyses suggest that Zld, Grh and Twi pioneer at regions within naïve or repressed chromatin, and that the inability of these factors to promote chromatin accessibility at class II regions is unlikely to be due to differences in preexisting features of chromatin.

### Chromatin opening by pioneer factors is driven by motif content

To further determine the features that drive the different outcomes on chromatin accessibility at PF-bound regions, we analyzed enrichment of the canonical Zld, Grh or Twi motifs for each of the three bound classes. We calculated the percentage of class I, II or III peaks that contained an instance of the canonical motif for the respective factor (Fig. 3c-e). When compared to all accessible regions, class I regions had low levels of enrichment for the canonical motifs, supporting our prior analysis suggesting that binding to these regions is nonspecific. By contrast, class II and III sites were highly enriched for the canonical motifs (Fig. 3c-e). Class III sites had the highest percentage of sites containing a canonical motif. We hypothesized that differences in motif content could explain why binding to class II regions does not result in chromatin opening. We therefore analyzed the number of motifs per peak and motif strength (when scored against the canonical motif) for class I-III regions. For Grh and Twi, but not for Zld, class III sites had more motifs per peak than class II sites (Fig. 3f,h,j). For all three factors, class III sites tended to have stronger motifs than class II sites (Fig. 3g,i,k). These data suggest that more and stronger motifs at class III as compared to class II sites facilitates the ability of Zld, Grh, and Twi to open chromatin.

Previous work showed that binding of some PFs depends on how target motifs are positioned on nucleosomes^18,56^. To test if pioneering by Zld, Grh or Twi was associated with motif positioning on nucleosomes, we analyzed published MNase-seq data from S2 cells^57^. Analysis of average MNase-seq signal at class I-III peaks indicated that class III sites have high levels of nucleosome occupancy that, on average, occur over the respective motif (Extended Data Fig. 6a-c). To test if there were different patterns of nucleosome occupancy at individual sites that were obscured by global analysis, we performed hierarchical clustering on the MNase-seq data. This analysis revealed that there was no preference for binding or chromatin opening at sites that had motifs in a particular orientation on nucleosomes (Extended Data Fig. 6d-f). For class II and III peaks, some motifs were positioned close to the nucleosome dyad, some motifs were at the edge of nucleosomes, and some motifs were in linker DNA. Additionally, the MNase signal in many regions did not show strongly positioned nucleosomes, likely because nucleosomes at silent regions often are not strongly positioned^58,59^. While some PFs may recognize their motif when specifically positioned on a nucleosome, Zld, Grh and Twi do not have a strong preference for motif positioning when exogenously expressed.

### Cell-type specific variables affect Zld, Grh and Twi binding

Prior studies have extensively characterized the *in vivo* binding sites of Zld, Grh, and Twi during normal *Drosophila* development. To better understand how cell-type specific variables affect PF binding, we compared ChIP-seq data for our PFs expressed in culture to published datasets for endogenous Zld, Grh and Twi binding. We analyzed Zld ChIP-seq data from the early embryo (nuclear cycle 14 embryos)^60^ and from larval neural stem cells^30^, Grh ChIP-seq data from late embryos (16-17 hours) and wing imaginal discs^31^ and Twi ChIP-seq data from early embryos (1-3 hours)^61^. We identified many binding sites that were bound *in vivo* but were not bound upon ectopic expression in S2 cells (Extended Data Fig. 7a-c). Chromatin enriched for repressive histone modifications H3K9me3 and H3K27me3 is resistant to binding by some PFs^6,12^. To test if these repressive marks were present in S2 cells at the resistant sites (bound *in vivo* but not upon ectopic expression), we compared our PF-binding data to published S2 cell ChIP-seq datasets for H3K27me3^51^ and H3K9me3^52^. We divided the sites bound *in vivo* but not in our S2 cell system into three classes: class IV sites have high levels of H3K27me3, class V sites have high levels of H3K9me3, and class VI have low levels of both repressive marks (Fig 4a-c; Extended Data Fig. 7d-l). Despite the presence of class IV, V and VI binding sites for all three proteins, there were distinct patterns of binding across different tissues. Zld binding in S2 cells was more similar to that of neural stem cells (NSC) than embryos, with many of the strongest binding sites in embryos remaining unbound in S2 cells (Fig. 4a). This differed for Grh, in which many of the strongest binding sites were shared across the three tissues (Fig. 4b). Twi binding was largely distinct in S2 cells and the embryo, with only a small subset of binding sites shared across the two tissues (Fig. 4c; Extended Data Fig. 7c).

**Figure 4:**
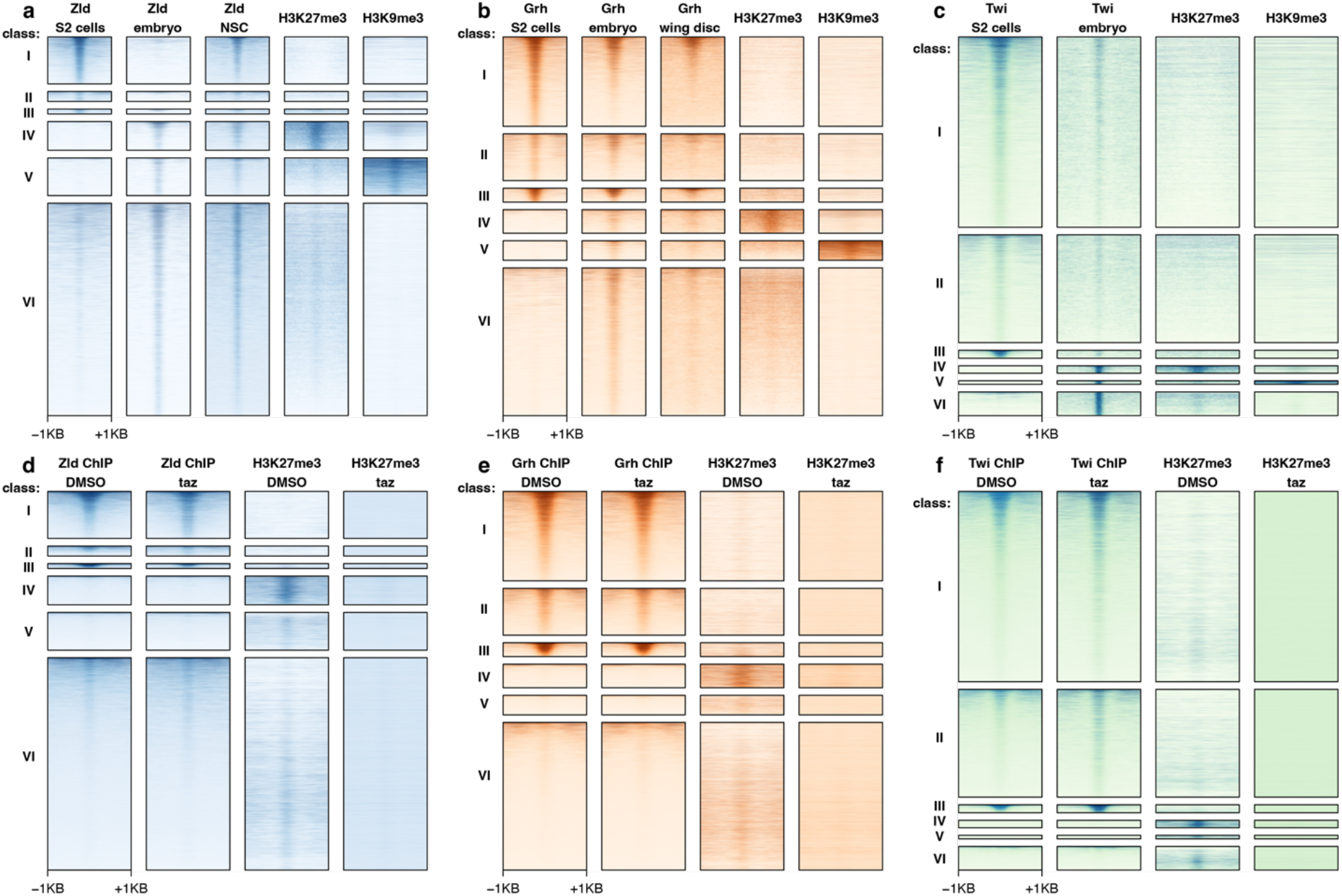
Many endogenous binding sites are resistant to ectopic pioneer-factor binding. **a-c,** Heatmaps comparing ChIP-seq signal in across different tissues for Zld **(a)**, Grh **(b)**, or Twi **(c)**. ChIP-seq data for repressive histone modifications H3K27me3^51^ and H3K9me3^52^ in wild-type S2 cells are also shown. **d-f,** Heatmaps comparing DMSO or tazemetostat (taz)-treated cells for Zld **(d)**, Grh **(e)**, or Twi **(f)**. Zld, Grh, and Twi ChIP-seq data are z-score normalized. CUT&RUN data for H3K27me3 is spike-in normalized using barcoded nucleosomes (see methods).

To test the functional relevance of repressive chromatin to PF binding, we used the chemical inhibitor tazemetostat to disrupt activity of the PRC2 complex responsible for depositing H3K27me3 (Extended Data Fig 8). Despite causing dramatic reductions in H3K27me3 levels, tazemetostat treatment did not result in global changes to Zld, Grh, or Twi binding when compared to DMSO controls (Fig. 4d-f). The majority of class IV binding sites remained unbound in tazemetostat-treated cells, demonstrating that loss of PRC2 activity is insufficient to promote widespread binding of any of the three factors studied.

### Pioneer-factor binding and chromatin opening is concentration dependent

Previous studies proposed that PF activity depends on protein concentration^62,63^. Therefore, we tested if differences in the protein concentration of Zld, Grh, or Twi could affect binding and pioneering. We performed ChIP-seq on cells induced to express Zld, Grh and Twi at a range of protein concentrations: uninduced, subphysiological, physiological, or supraphysiological (Extended Data Fig. 9a-c; see methods). Analysis of our previously defined class I-VI sites showed that binding was concentration dependent (Fig. 5). At class II and III binding sites, subphysiological levels of Zld, Grh, and Twi were sufficient for some, albeit reduced binding (Fig. 5a,e,i). Expression of supraphysiological protein levels resulted in increased ChIP intensity at class II and III sites (Fig. 5a,e,i). To identify the relationship between PF binding and chromatin accessibility, we performed ATAC-seq on cells with the same range in expression. As we identified for PF binding, changes in chromatin accessibility were concentration dependent, with more limited chromatin opening at lower concentrations and more robust opening at higher concentrations (Fig. 5b,f,j). Many class II binding sites became accessible when Zld, Grh, and Twi were expressed at the highest levels, suggesting that increased chromatin occupancy, as assayed by ChIP-seq, could result in chromatin accessibility. For Grh and Zld, but not Twi, supraphysiological concentrations allowed some binding and opening to class IV-VI sites, which were not bound at lower concentrations (Extended Data Fig. 9d-f).

**Figure 5:**
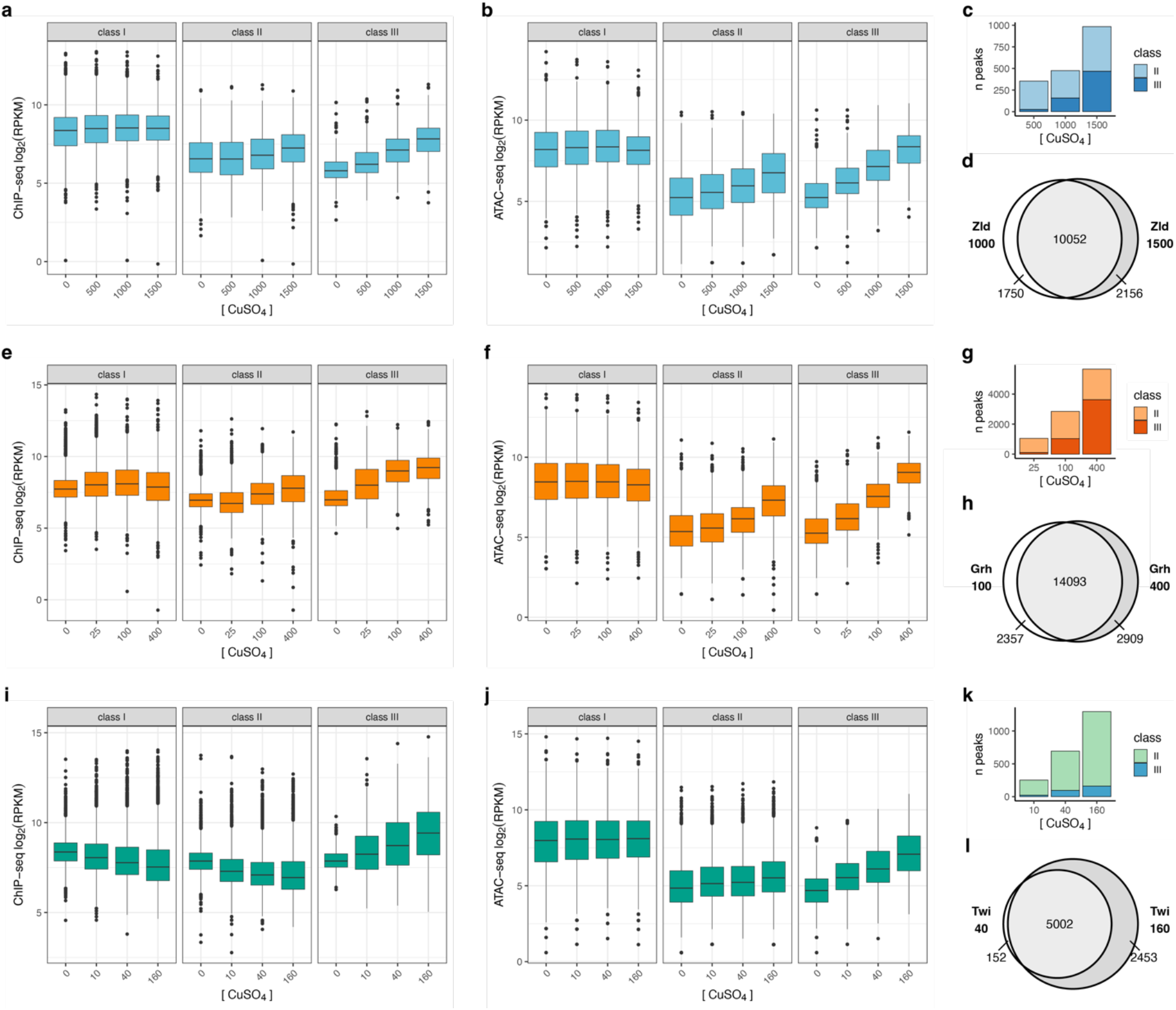
Pioneer-factor binding and opening of closed chromatin is concentration dependent. **a,e,i,** Boxplots of ChIP-seq signal at class I-III regions for Zld **(a)**, Grh **(e)**, or Twi **(i)** when expressed at different levels. **b,f,j,** Boxplots of ATAC-seq signal at class I-III regions for Zld **(b)**, Grh **(f)**, or Twi **(j)** when expressed at different levels. **c,g,k,** Distribution of class II and III peaks when defined using ChIP-seq and ATAC-seq data for each concentration of CuSO_4_ for Zld **(c)**, Grh **(g)**, or Twi **(k)**. Y-axis indicates the total number of peaks called at each concentration. Colors indicate the proportion of those peaks that are defined as class II, or III. **d,h,l,** Venn diagrams showing overlap of peaks called at physiological or supraphysiological expression levels for Zld **(d)**, Grh **(h)**, or Twi **(l)**.

To determine how protein concentration changed the distribution of class I, II and III sites, we redefined these classes using the ChIP-seq and ATAC-seq data at each concentration. For all three factors, supraphysiological concentrations led to an increase in binding to closed chromatin (Fig. 5c,g,k). For Twi, this increase was mostly within class II regions. For Grh and Zld, supraphysiological concentrations caused a dramatic increase in class III regions relative to class II (Fig. 5c,g). To assess the extent to which increased concentrations led to binding to new regions that were not bound at physiological concentrations, we overlapped the binding sites detected at physiological vs. supraphysiological concentrations (Fig. 5d,h,l). For all three proteins, there were novel binding sites that were bound only when expressed at the highest concentration. Together, these data demonstrate that both binding and chromatin opening by PFs are concentration dependent, and that increasing protein concentration leads to novel binding sites, including some endogenous binding sites for these factors during normal development.

### Regions outside the DNA-binding domain are required for binding and opening of closed chromatin

Binding to nucleosomes by PFs requires recognition of target motifs by DNA-binding domains (DBDs)^4^. However, little is known about how regions outside the DBD may contribute to pioneering function. We generated stable cell lines expressing the Zld or Grh DBD at levels comparable to the full-length protein (Extended Data Fig. 10a,b). Immunostaining confirmed that the each DBD alone was properly localized to the nucleus (Extended Data Fig. 10c-d). We performed ChIP-seq and ATAC-seq on cells expressing approximately physiological levels of the DBDs alone. Analysis of ChIP-seq signal at our previously defined class I-III regions revealed that the Zld DBD bound robustly to class I regions, but was unable to bind closed chromatin at class II or III regions (Fig. 6a,c). Binding of the Grh DBD to class II-III regions was strongly reduced compared to the full-length protein, although some closed regions retained low levels of binding (Fig. 6b,e). For both Zld and Grh, the DBD alone was insufficient to open chromatin (Fig 6d,f). Together, these experiments show that regions outside the DBD are essential for binding and opening closed chromatin.

**Figure 6:**
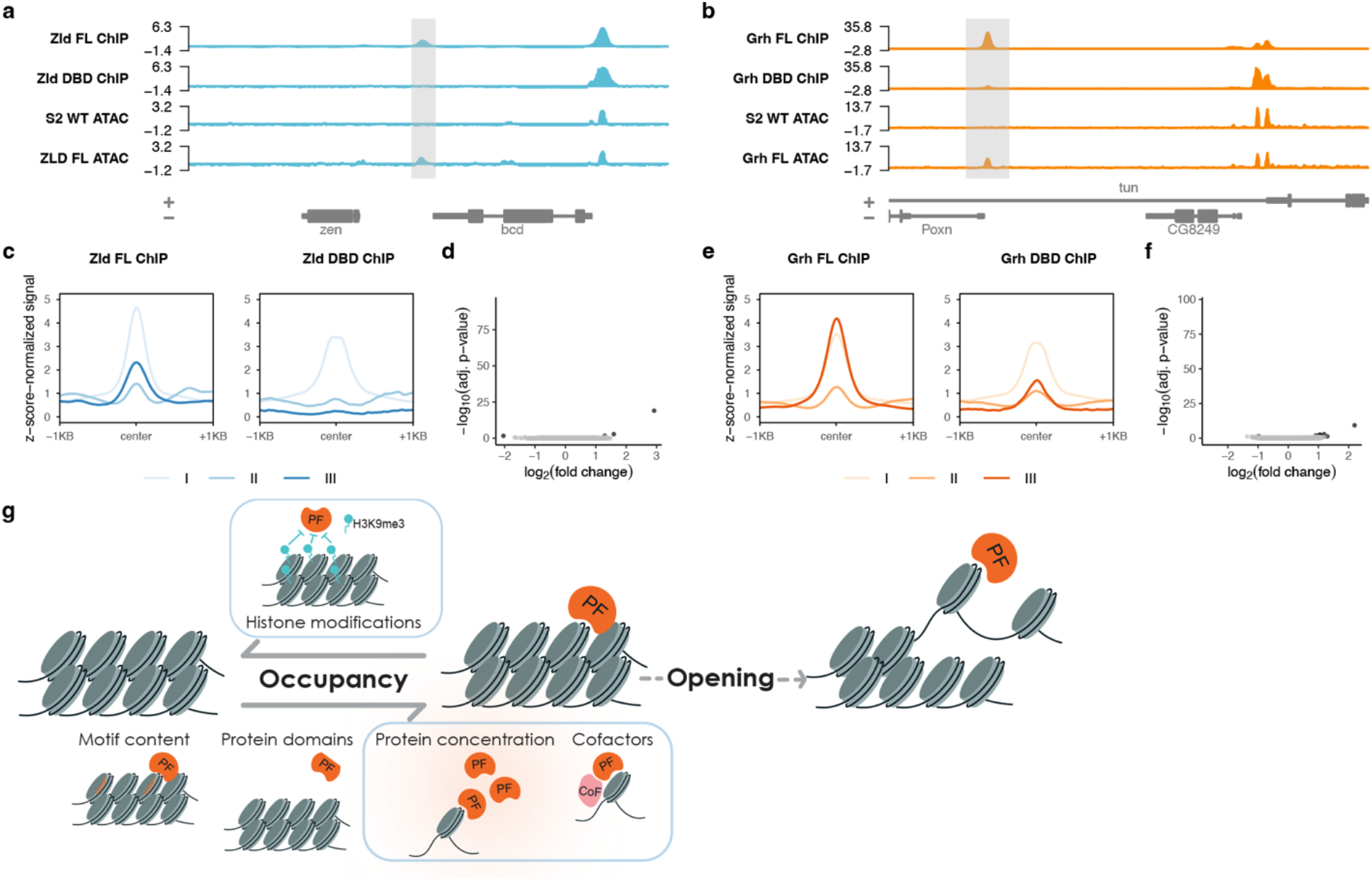
Regions outside the DNA-binding domain are required for pioneer-factor function. **a,b,** Genome browser tracks showing example of class III regions where expression of the DBD alone is insufficient for robust binding and chromatin opening. Adjacent class I regions are shown where the DBD alone can bind. Tracks shown for Zld **(a)** and Grh **(b)**. Class III regions are highlighted with gray shading. **c,e,** Metaplots comparing ChIP-seq signal for full-length protein or DBD alone at class I, II and III regions for Zld **(c)** or Grh **(e)**. **d,f,** Volcano plots for ATAC-seq data showing the absence of changes to chromatin accessibility upon expression of Zld **(d)** or Grh DBD **(f)**. **g,** Model for the regulation of PFs. A high level of chromatin occupancy is required for PFs to initiate chromatin opening. Chromatin occupancy can be regulated by protein-intrinsic features such as motif affinity and protein domains, and by cell-intrinsic properties such as histone modifications, PF concentration, and cell-type specific cofactors (outlined in blue).

## Discussion

To identify the mechanisms that regulate tissue-specific PF engagement with chromatin, we exogenously expressed Zld, Grh, and Twi in S2 cells where they are not normally expressed. Leveraging prior data identifying binding sites for these factors *in vivo* and analyzing the wealth of data on chromatin structure in S2 cells, we identified features that govern PF activity. We studied the two well-defined PFs Zld and Grh, along with Twi, which possess features distinct from these two factors. Studying these three distinct TFs, which engage the genome through structurally distinct DBDs, allowed us to determine shared properties that regulate their pioneering activity. Furthermore, our genome-wide analysis of binding, accessibility and gene expression allowed us to separate PF occupancy and activity. Collectively, we find that pioneering activity requires a high level of chromatin occupancy that can be achieved through both protein-intrinsic and protein-extrinsic features such as local chromatin structure, protein concentration, DNA motif content, and protein domains outside the DBD (Fig. 6g).

Even when expressed at low levels Zld, Grh and Twi bind promiscuously to active, accessible regions with degenerate motifs or no detectable motifs at all. When expressed at approximately physiological levels these factors all possess the capacity to bind to canonical motifs in closed chromatin. However, these factors are largely excluded from repressed chromatin enriched for H3K9me3 or H3K27me3. This binding to naïve chromatin is similar to what has previously been reported for other pioneer factors^6,7^. Regions with high levels of H3K9me3 are resistant to binding by the reprogramming factors Oct4, Sox2 and Klf4, and c-Myc (OSKM). Depletion of H3K9 methyltransferases increases OSKM occupancy at these regions, suggesting that this repressive chromatin state is a barrier to PF binding^6^. Similarly, cell-type specific binding of the PF Pax7 is anti-correlated with levels of H3K27me3^64^. Here, we specifically tested whether H3K27me3 regulates cell-type specific occupancy of Zld, Grh and Twi using the PRC2 inhibitor tazmetostat. While H3K27me3 levels were decreased in our treatment, PF binding was largely unchanged. Thus, unlike H3K9me3 in the case of OSKM, H3K27me3 is not a barrier to binding by the three factors assayed and suggest other features are restricting binding to these regions.

While depleting H3K27me3 levels did not result in Zld, Grh and Twi binding to additional regions, increased expression of both Zld and Grh resulted in binding to regions that were unbound at lower expression levels. These results suggest that protein concentration and affinity for the target motif drives tissue-specific PF binding rather than the presence or absence of Polycomb-mediated silencing. Tissue-specific PF occupancy is also likely regulated by the collection of cofactors expressed in a given tissue. This may partially explain the presence of class VI sites that lack repressive histone modifications in S2 cells and are bound *in vivo* but not when expressed exogenously. For example, in mammalian cell culture FOXA2 occupancy at a subset of tissue-specific binding sites was increased when GATA4 was present^7^. Similarly, PARP-1 stabilizes Sox2 binding at subset of physiologically relevant sites in mouse embryonic stem cells^65^. Unlike Zld and Grh, increased expression of Twi did not result in binding to regions bound in other tissues. It is possible that additional cofactors may be required for Twi occupancy at these regions. Along with prior studies, our data suggest that PF occupancy is regulated by tissue-intrinsic features, including levels of PF expression, the complement of cofactors expressed and chromatin structure.

In addition to cell-type specific factors, we demonstrate that protein-intrinsic features control PF binding. We show that regions outside the DBD of Zld and Grh are important for binding-site selection. Similarly, regions outside the DBDs of human PFs FOXA1 and SOX2 are required for robust binding of closed chromatin^66^. Increasing evidence suggests that eukaryotic transcription factors are not as modular as their bacterial counterparts and has demonstrated a role for intrinsically disordered regions (IDRs) in regulating chromatin occupancy^67,68^ Both Zld and Grh contain long disordered domains. For Zld, this disordered domain is required for transcriptional activation and promotes optimal nucleosome binding *in vitro*^24,69^. IDRs may contribute to TF binding through protein-protein interactions with other TFs, nonspecific interactions with DNA or histones, or by driving condensate formation^70,71^. Indeed, Zld is visualized in hubs within the nucleus, and these promote binding of additional transcription factors^72,73^. Zld and Grh DBDs retain the capacity to bind to class I regions, despite the weaker motifs present at these sites. Thus, the binding to class I regions is not due to interactions between IDRs and other proteins present at active chromatin. Instead, this binding may be driven by the high affinity of the DBDs for nucleosome-free DNA. To provide a more complete understanding of how cis-regulatory regions are established, we must further investigate how IDRs contribute to PF binding and chromatin accessibility.

The failure of the DBD alone to promote chromatin accessibility implicates a separation between the ability to bind nucleosomes and the capacity to promote chromatin accessibility. While the DBD of Zld is sufficient to bind nucleosomes *in vitro*, regions outside the DBD promote this interaction^24^. Thus, nucleosome binding *in vitro* does not necessarily translate into *in vivo* occupancy of closed chromatin. Local restructuring of nucleosome-DNA interactions may be insufficient for establishing accessible chromatin^18,74^ and, instead, may stabilize PF binding to allow recruitment of cofactors, such as chromatin remodelers^16,17^. Furthermore, our identification of hundreds of class II regions, which are bound but not opened by exogenous PF expression, distinguishes chromatin binding and PF activity. We propose that PF-mediated chromatin accessibility requires high chromatin occupancy. This is supported by our data demonstrating a higher motif content at sites in which PF binding promotes accessibility as compared to sites which remain closed despite PF occupancy. Furthermore, we identified a strong concentration dependence of Zld and Grh for chromatin opening. This model would suggest that PF expression levels must be tightly controlled during development. Indeed, misexpression of the PFs Grh and DUX4 leads to diseases such as epithelial cancer or facioscapulohumeral muscular dystrophy (FSHD), respectively^75,76^. Together our data identify protein intrinsic and extrinsic features that govern PF binding and activity, providing insights into how these factors define novel *cis*-regulatory elements and drive gene-regulatory networks to modulate cell fate. A deeper understanding of how PF activity is regulated will have important implications for determining how diseases are caused by their misexpression and our ability to use these powerful factors for cellular reprogramming.

## Materials and Methods

### Cell culture and generation of stable cell lines

*zld*, *grh*, or *twi* cDNA (isoforms RB, RH, and RA, respectively) were cloned into pMT-puro plasmid (Addgene) via Gibson cloning (New England Biolabs). For cloning of the Zld and Grh DNA-binding domains, DNA encoding amino acids 1114-1487 (Zld) or amino acids 603-1032 (Grh) was cloned into pMT-puro.

We validated our wild-type S2 cells by performing RNA-seq and verifying the expression of known S2 cell markers and the absence of Kc cell markers^77^. S2 cells were cultured at 27°C in Schneider’s medium (Thermo Fisher Scientific) supplemented with 10% fetal bovine serum (Omega Scientific) and antibiotic/antimycotic (Thermo Fisher Scientific). For the generation of stable cell lines, cells were plated at 5×10^5^ cells per mL. After 24 hours, cells were transfected with 10 µg plasmid DNA using Effectene transfection reagent (Qiagen). After an additional 24 hours, puromycin was added to a final concentration of 2 µg/mL. Stable cell lines were recovered after 2-3 weeks of selection. Following recovery of stable cell lines, cells were cultured with 1 µg/mL puromycin.

### Induction of transcription factor expression

Transcription factor expression was induced by adding CuSO_4_ to the cell culture media. Expression of protein at physiological levels was achieved with 1000, 100, or 40 µM CuSO_4_ for Zld, Grh or Twi, respectively (Extended Data Fig. 1c-d; Extended Data Fig. 4a). For experiments with sub- or supraphysiological protein levels (Fig. 5; Extended Data Fig. 9a-c), the following CuSO_4_ concentrations were used: 500 µM (Zld subphysiological), 1000 µM (Zld physiological) or 1500 µM (Zld supraphysiological); 25 µM (Grh subphysiological), 100 µM (Grh physiological) or 400 µM (Grh supraphysiological); 10 µM (Twi subphysiological), 40 µM (Twi physiological) or 160 µM (Twi supraphysiological). The Zld or Grh DNA-binding domains were induced with 100 µM, or 20 µM CuSO_4_, respectively, for ChIP, ATAC and immunofluorescence experiments (Extended Data Fig. 10). For all induction experiments, cells were plated at 1×10^6^ cells per mL. Unless otherwise specified, experiments were performed with cells harvested 48 hours after induction.

### Immunoblotting

Proteins were separated on denaturing polyacramide gels before transfer to a 0.45 μm Immobilon-P PVDF membrane (Millipore) in transfer buffer (25 mM Tris, 200 mM Glycine, 20% methanol) for 60 min (75 min for Zld) at 500mA at 4°C. The membranes were blocked with BLOTTO (2.5% non-fat dry milk, 0.5% BSA, 0.5% NP-40, in TBST) for 30 min at room temperature and then incubated with anti-Zld (1:750)^78^, anti-Grh (1:1000)^78^, anti-Twi (1:1000)^79^, anti-HA-peroxidase (1:500) (clone 3F10, Roche), or anti-Tubulin (DM1A, 1:5000) (Sigma), overnight at 4°C. The secondary incubation was performed with goat anti-rabbit IgG-HRP conjugate (1:3000) or goat anti-mouse IgG-HRP conjugate (1:3000) (Bio-Rad) for 1 hr at room temperature. Blots were treated with SuperSignal West Pico PLUS chemiluminescent substrate (Thermo Fisher Scientific) and visualized using the Azure Biosystems c600 or Kodak/Carestream BioMax Film (VWR).

### Immunostaining

Immunostaining was performed as described previously^80^. Briefly, 1×10^6^ cells in 200 µL PBS were applied to a coverslip pre-coated with 10% poly-L-lysine. Cells were fixed with 4% methanol-free formaldehyde for 30 minutes at 37°C, permeabilized with 0.2% Triton-X 100 for 10 minutes at room temperature, and blocked with 1% BSA for 1 hour. Cells were incubated with a 1:500 dilution of anti-Zld or anti-Grh primary antibody overnight at 4°C. After washing with PBS, cells were incubated with a 1:500 dilution of goat anti-rabbit IgG DyLight 488 conjugated secondary antibody (Thermo Fisher Scientific #35552) for 1 hour at RT. Imaging was performed on a Nikon Ti2 epi-fluorescent microscope with a 60x objective lens.

### ChIP-seq

For each ChIP experiment, ∼50 million cells were crosslinked by adding methanol-free formaldehyde directly to the cell culture medium to a final concentration of 0.8%. Cells were rotated on a nutator for 7 minutes at room temperature before quenching of crosslinking by adding glycine to 125 mM. Crosslinked cells were pelleted by centrifugation at 600 × *g* for 3 minutes. Cells were washed twice with 1X PBS. Cells were lysed by resuspending in lysis buffer (50 mM HEPES pH 7.9, 140 mM NaCl, 1 mM EDTA, 10% glycerol, 0.5% NP40, 0.25% Triton-X 100) with protease inhibitors (Pierce) and incubating on ice for 10 minutes. The resulting chromatin was pelleted for 5 minutes at 1500 × *g* and resuspended in RIPA buffer. Chromatin was sonicated in a Covaris S220 Ultrasonicator (5 cycles of 120 seconds with 60 second delay, 170 peak power, 10% duty factor, 200 cycles/burst). Sonicated chromatin was centrifuged for 10 minutes at 10,000 × *g* to pellet insoluble material. 5% of the supernatant was set aside as input, and the remainder was incubated at 4°C overnight with antibodies (6 µL anti-Zld, 8 µL anti-Grh, 10 µg anti-Twi, 7.5 µL anti-HA). 20 µL protein A beads (Dynabeads Protein A, ThermoFisher Scientific) blocked with BSA were added and samples were incubated at 4°C for 4 hours. Beads were separated on a magnet and washed 3× with low salt wash buffer (10 mM Tris pH 7.6, 1 mM EDTA, 0.1% SDS, 0.1% Na-Deoxycholate, 1% Triton-X 100, 150 mM NaCl), 2× high salt wash buffer (10 mM Tris pH 7.6, 1 mM EDTA, 0.1% SDS, 0.1% sodium deoxycholate, 1% Triton-X 100, 300 mM NaC), 2× LiCl wash buffer (0.25 M LiCl, 0.5% NP40, 0.5% sodium deoxycholate), and 1× with TE buffer with NaCl (10 mM Tris-HCl pH 8.0, 1 mM EDTA, 50 mM NaCl). Beads were resuspended in elution buffer (50 mM Tris-HCl pH 8.0, 1% SDS, 10 mM EDTA) and incubated at 65°C for 10 minutes. The IP and input samples were treated with 4.5 µL RNAse A for 30 minutes. 5 µL Proteinase K was added and samples were incubated overnight at 65°C to reverse crosslinking. DNA was isolated by phenol:chloroform extraction, precipitated with EtOH, and resuspended in 20 µL H_2_O.Preparation of sequencing libraries was performed using NEBNext Ultra II kit (NEB) with 7 PCR cycles for library amplification. Sequencing was performed on the Illumina Hi-Seq4000 using 50bp single-end reads, or on the Illumina NovaSeq 6000 using 150bp paired-end reads.

### ChIP-seq analysis

Bowtie2 version 2.4.4^81^ was used to align ChIP-seq reads to the *Drosophila melanogaster* genome (version dm6) with the following non-default parameters: `-k 2 --very-sensitive --no-mixed --no- discordant -X 5000`. Aligned reads were filtered to include only reads with a mapping quality score > 30. Reads mapping to unplaced scaffolds or the mitochondrial genome were discarded. Peak calling was performed using MACS2 version 2.2 with parameters `-g dm --call-summits`. Data from ChIP experiments performed with the same antibody in wild-type cells were used as a control for peak calling. Only peaks that were detected in all replicates were considered in downstream analysis. bigWig files were generated using deepTools bamCoverage version 3.5.1 with parameters `–binSize 10`. bigWig files were z-score normalized by subtracting the genome-wide average of all bins from each bin value and dividing by the standard deviation of all bins.

### RNA-seq

5×10^6^ cells were harvested and resuspended in 800 µL Trizol (Invitrogen). RNA was purified using a chloroform extraction followed by a column-based Quick-RNA MiniPrep extraction (Zymo). Poly-A-selected RNA sequencing (RNA-seq) libraries were prepared using the TruSeq RNA sample prep kit v2 (Illumina).

### RNA-seq analysis

Raw reads were aligned to the *Drosophila melanogaster* (dm6) genome using HISAT2 version 2.1.0 with parameters `-k 2 –very-sensitive`. Reads with a mapping quality score < 30 and reads aligning to the mitochondrial genome or unplaced scaffolds were discarded. FeatureCounts (from Subread version 2.0) was used to assign aligned reads to genes (FlyBase gene annotation release r6.45). The resultant table of read counts was used to perform differential expression analysis using DESeq2^36^. Genes with an adjusted p-value < 0.05 and a fold change > 2 were considered differentially expressed.

### ATAC-seq

ATAC-seq was performed as described previously^82^. 2×10^5^ cells were harvested for ATAC-seq by centrifugation at 600 × *g* for 3 minutes at RT. Cells were washed once in 100 µL 1X PBS and resuspended in 100 µL ATAC lysis buffer (10 mM Tris pH 7.5, 10 mM NaCl, 3 mM MgCl_2_,0.1% NP-40). Cells were centrifuged at 600 × *g* for 10 minutes at 4°C. Supernatant was removed and pellet was resuspended in 47.5 µL buffer TD (Illumina). 2.5 uL of Tagment DNA Enzyme (Illumina) was added and samples were incubated at 37°C for 30 minutes. Tagmented DNA was purified using MinElute Reaction Cleanup Kit (Qiagen) and eluted in 10 µL of buffer EB. DNA was PCR amplified for 12 cycles with the following conditions:72°C for 5 minutes, 98°C for 30 seconds, then 12 cycles of 98°C for 10 seconds, 63°C for 30 seconds and 72°C for 1 minute. Amplified libraries were purified using a 1.2X ratio of Axygen paramagnetic beads. Sequencing was performed on the Illumina NovaSeq 6000 using 150bp paired-end reads.

### ATAC-seq analysis

Raw ATAC-seq reads were trimmed to remove adapter sequences using NGmerge version 0.3^83^. Trimmed reads were aligned to the *Drosophila melanogaster* genome (version dm6) using bowtie2 with paramters `-k 2 --very-sensitive --no-mixed --no-discordant -X 5000`. Reads with a mapping quality < 30 and reads aligning to the mitochondrial genome or scaffolds were discarded. As described previously, only fragments < 100bp were considered for downstream analysis^82^. Reads were combined across all replicates and peak calling was performed using MACS2 with parameters `-f BAMPE --keep- dup all -g dm --call-summits`. featureCounts was used to count the number of reads aligning within 200bp (100bp upstream or downstream) of peak summits. Generation of z-score normalized bigWig files was performed as described above for ChIP-seq data. Differential accessibility analysis was performed using DESeq2. Regions with an adjusted p-value < 0.05 were considered differentially accessible.

### Treatment of S2 cells with tazemetostat

Cells were plated at 1×10^6^ cells per mL and DMSO or tazemetostat (Fisher Scientific) was added to a final concentration of 10 µM. Cells were split on days 2 and 5 and tazemetostat was added to maintain a final concentration of 10 µM. On day 5, CuSO_4_ was added to achieve physiological expression of Zld, Grh or Twi (see above). On day 7, cells were harvested for immunoblotting, ChIP and CUT&RUN.

### CUT&RUN

CUT&RUN was performed using EpiCypher reagents according to the manufacturer protocol. 2×10^5^ cells were used for each CUT&RUN reaction. Overnight incubation with antibodies was performed in antibody buffer containing 0.05% digitonin. 0.5 µL anti-H3K27me3 (Cell signaling technologies), anti-Zld, anti-Grh, or rabbit IgG were used for CUT&RUN. For H3K27me3 CUT&RUN, a 1:5 dilution of K-MetStat Panel spike-in nucleosomes (EpiCypher, cat. # 19-1002) was added to each reaction prior to addition of antibody. Libraries were prepared using the NEBNext Ultra II kit (NEB). During library preparation, cleanup steps were performed using a 1.1X ratio of Axygen paramagnetic beads, as recommended by EpiCypher. PCR amplification was performed with the following conditions: 98°C for 45 seconds, then 14 cycles of 98°C for 15 seconds, 60°C for 10 seconds, and a final extension of 72°C for 1 minute. Sequencing was performed on the Illumina NovaSeq 6000 with 150bp paired-end reads.

### CUT&RUN analysis

Trimming, alignment, and quality filtering of CUT&RUN reads was performed as described above for ATAC-seq data. Peak calling was performed using MACS2 with parameters `–broad -f BAMPE –keep- dup all -g dm` for H3K27me3 or parameters `-f BAMPE --keep-dup all -g dm --call-summits` for Zld and Grh. Generation of z-score normalized bigWig files was performed as described above for ChIP-seq data. For H3K27me3 data, the percentage of reads containing barcoded Epicypher nucleosomes was calculated and 1 divided by this percentage was used as the scaling factor for spike-in normalization. The z-score normalized read depth was multiplied by these scaling factors to generate spike-in normalized bigWigs.

### Analysis of previously published data

Previously published datasets analyzed in this study can be found in Supplementary Table 2. All previously published data were analyzed in parallel with data from this study using analysis parameters described above.

### Data and code availability

Sequencing data have been deposited in the Gene Expresion Omnibus under accession GSE227884. All analysis code is available on GitHub: https://github.com/tjgibson/S2_pioneers_manuscript.

## Acknowledgements

We thank Alex Theis, Meghan Freund, and Andrew Mehle for helpful discussions and advice. We thank Julia Zeitlinger for generously sharing the Twist antibody. We acknowledge the University of Wisconsin-Madison Biotechnology Center and the NUSeq Core Facility for sequencing. TJG was supported by the National Institutes of Health (NIH) National Research Service Award T32 GM007215. Experiments were supported by a R35 GM136298 from NIH to MMH. MMH was also supported by a Vallee Scholar Award. MMH is a Romnes Faculty Fellow and Vilas Faculty Mid-Career Investigator.

---

**Extended Data Figure 1:**
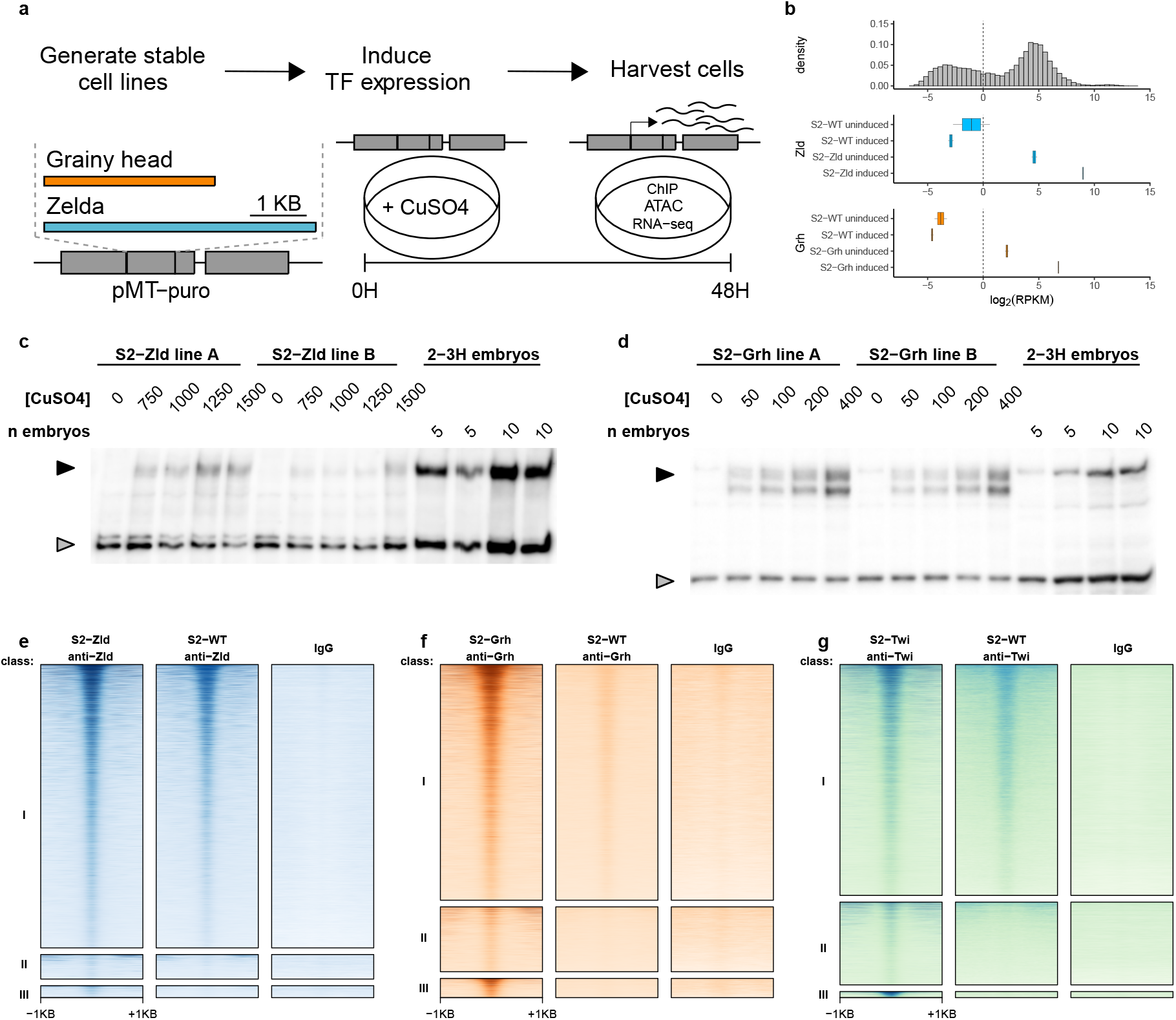
Stable cell lines allow inducible expression of transcription factors at physiological concentrations. **a,** Schematic of generation of stable cell lines and induction of protein expression. **b,** mRNA levels of *zld* and *grh* in S2 cells. Top histogram shows the distribution of mRNA levels for all *Drosophila* genes. Vertical dashed line indicates a log_2_ RPKM value of 0 as a threshold for considering a gene to be expressed. **c-d,** Immunoblots showing titration of Zld **(c)** or Grh **(d)** protein levels in stable cell lines. Two independently generated cell lines are shown and compared to 2-3 hours (H) embryos. 60,000 cells were loaded in each well, which is equivalent to the approximately 60,000 nuclei present in 10 2-3 hours embryos. Black arrowheads indicate Zld **(c)** or Grh **(d)**. Gray arrowheads indicate background bands used to assess loading. **e-g,** Heatmaps comparing Zld **(e)**, Grh **(f)**, or Twi **(g)** ChIP-seq signal to control experiments in which anti-Zld, anti-Grh, anti-Twi, or IgG antibodies were used to perform immunopreciptation in wild-type (WT) cells.

**Extended Data Figure 2:**
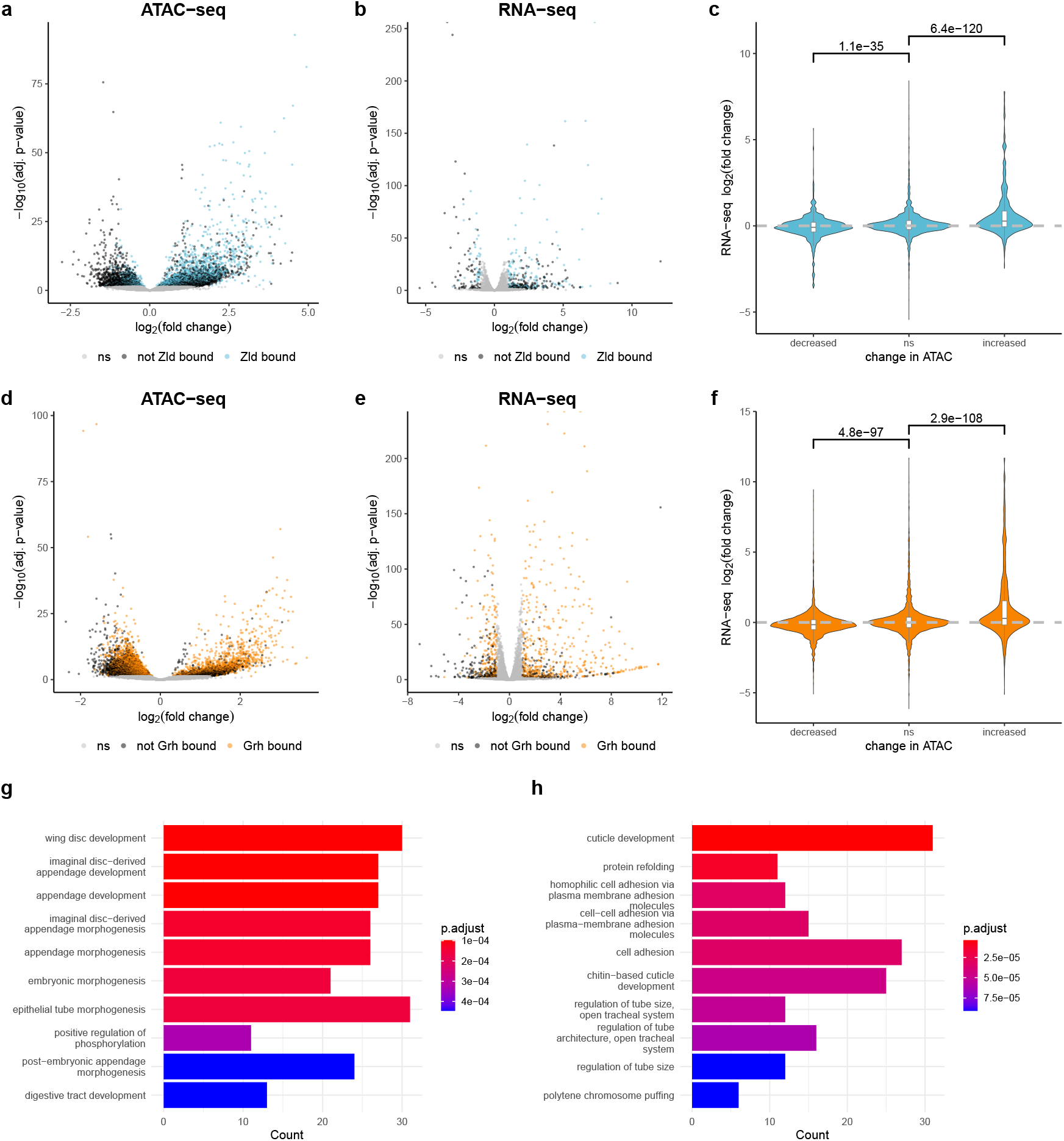
Ectopic expression of pioneer factors in S2 cells leads to widespread changes to chromatin accessibility and gene expression. **a,d,** Volcano plots showing changes in ATAC-seq signal in cells expressing Zld **(a)** or Grh **(d)** when compared to wild-type cells treated with the same concentration of CuSO_4_. **b,e,** RNA-seq volcano plots showing gene expression changes in cells expressing Zld **(b)** or Grh **(e)** when compared to wild-type cells treated with the same concentration of CuSO_4_. **c,f,** Violin plots showing the correlation between changes in chromatin accessibility and gene expression upon expression of Zld **(c)** or Grh **(f)**. On the x-axis, all ATAC-seq peaks are grouped based on increased, decreased, or non-significant (ns) changes to chromatin accessibility in Zld- or Grh-expresing cells compared to wild-type cells. Groups were compared using a Wilcoxon rank sum test and Bonferroni-corrected p-values are shown. **g,h,** Bar plots showing enrichment of gene ontology terms in genes significantly upregulated upon expression of Zld **(g)** or Grh **(h)**.

**Extended Data Figure 3:**
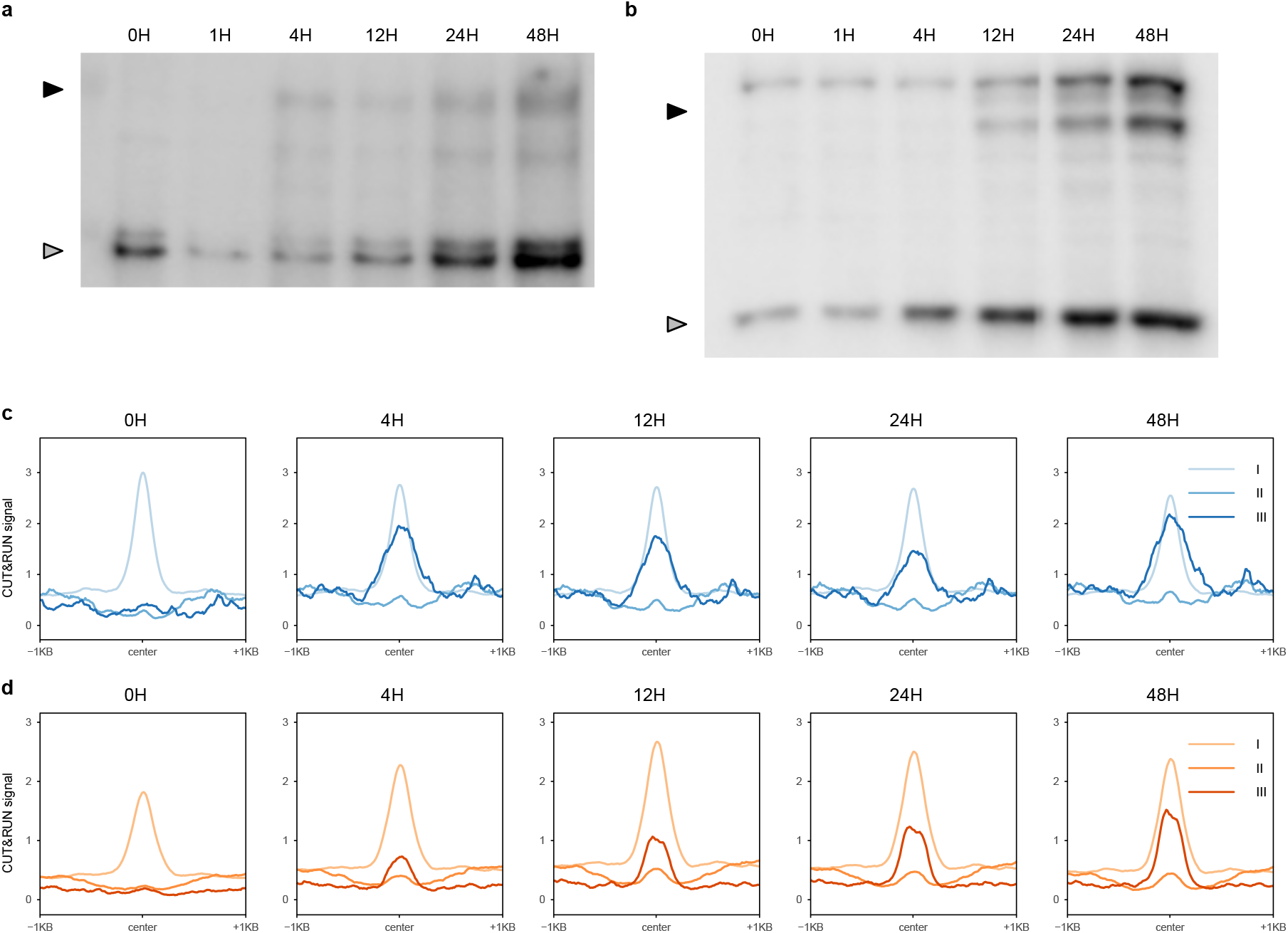
Zld and Grh bind to chromatin rapidly after induction of protein expression. **a-b,** Immunoblots showing time course of Zld **(a)** or Grh **(b)** protein expression following induction of stable cell lines. Black arrowheads indicate Zld **(a)** or Grh **(b)**. Gray arrowheads indicate background bands used to assess loading. **c-d,** Metaplots showing average z-score normalized CUT&RUN signal at class I, II or III sites at different time points following induction.

**Extended Data Figure 4:**
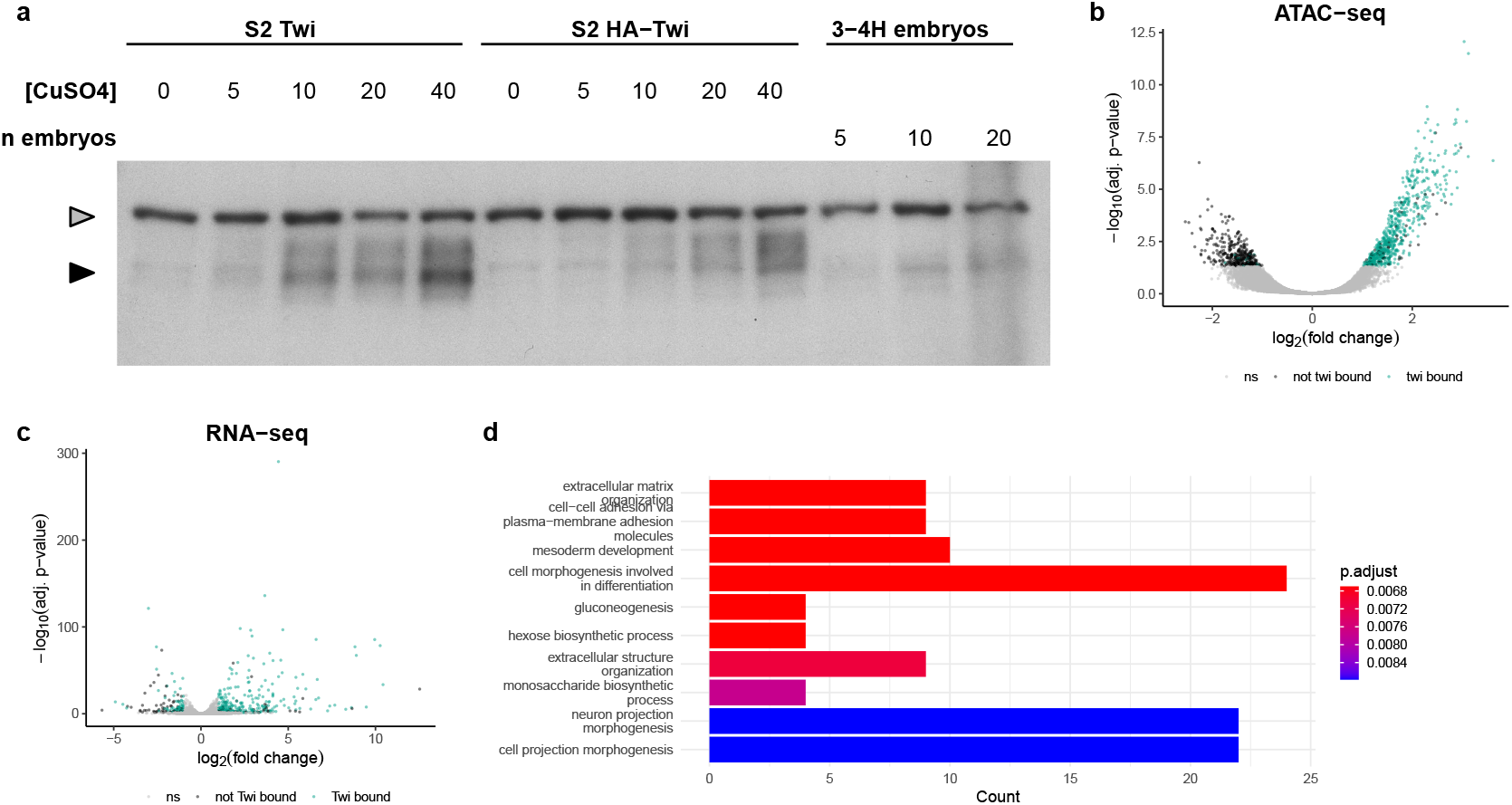
Twist binding leads to chromatin opening and transcriptional activation. **a,** Immunoblot showing titration of Twi or HA-Twi protein levels in stable cell lines. Protein levels in stable cell lines are compared to 3-4 hour (H) old embryos. Black arrowhead indicates Twi or HA-Twi. Gray arrowhead indicates background band used to assess loading. **b,** Volcano plots showing changes in ATAC-seq signal in cells expressing Twi when compared to wild-type cells treated with the same concentration of CuSO_4_. **c,** RNA-seq volcano plots showing gene expression changes in cells expressing Twi when compared to wild-type cells treated with the same concentration of CuSO_4_. **d,** Bar plots showing enrichment of gene ontology terms in genes significantly upregulated upon expression of Twi.

**Extended Data Figure 5:**
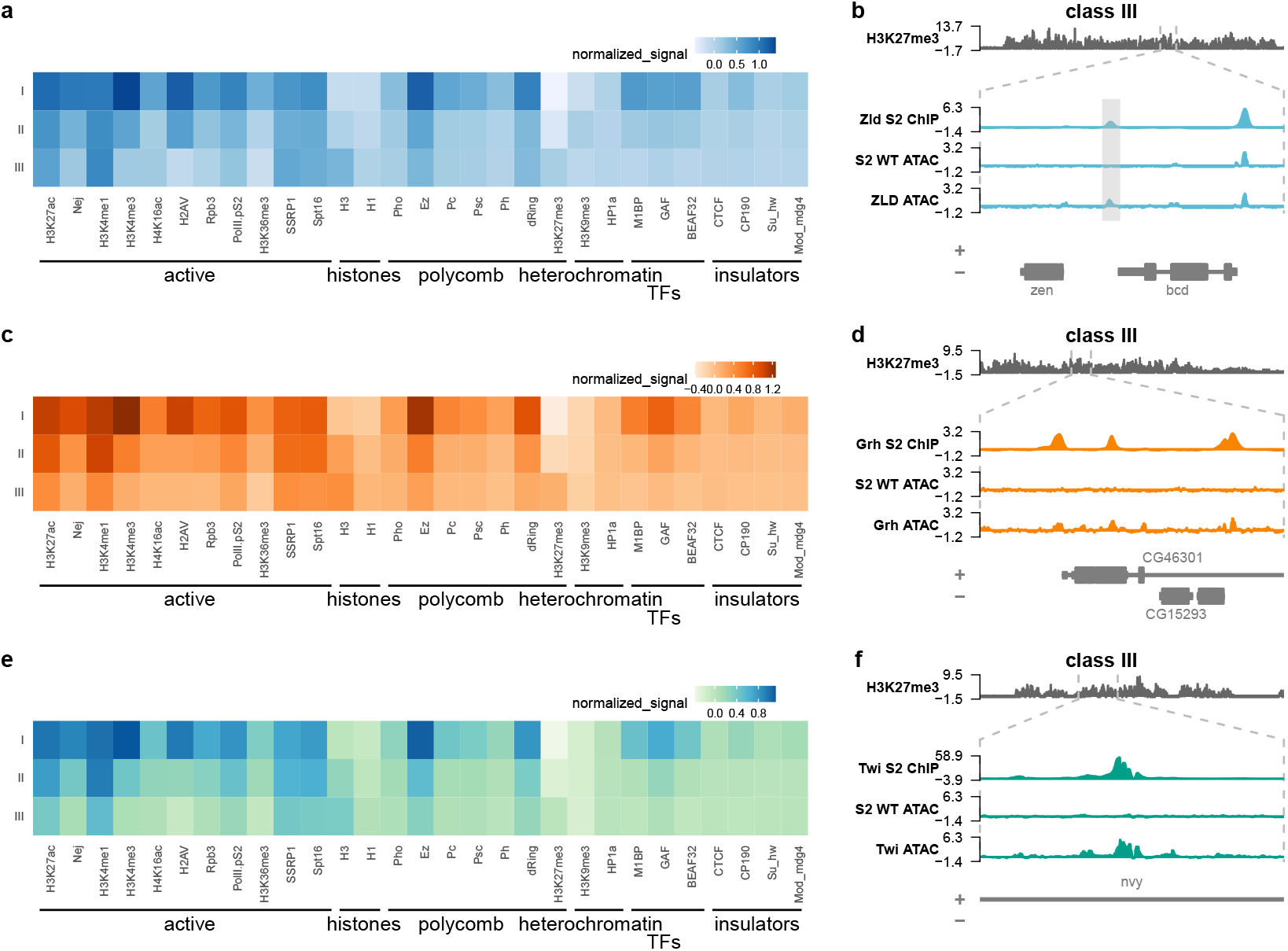
Chromatin features associated with Zld, Grh and Twi binding sites. **a,c,e,** Heatmaps showing the levels of different chromatin marks in class I, II or III regions for Zld **(a)**, Grh **(c)** or Twi **(e)**. The color represents the average z-score normalized read depth across a 1 KB region surrounding the center of class I, II or III ChIP-seq peaks. **b,d,f,** Example genome browser tracks for class III regions with high levels of H3K27me3 for Zld **(b)**, Grh **(d)**, or Twi **(f)**.

**Extended Data Figure 6:**
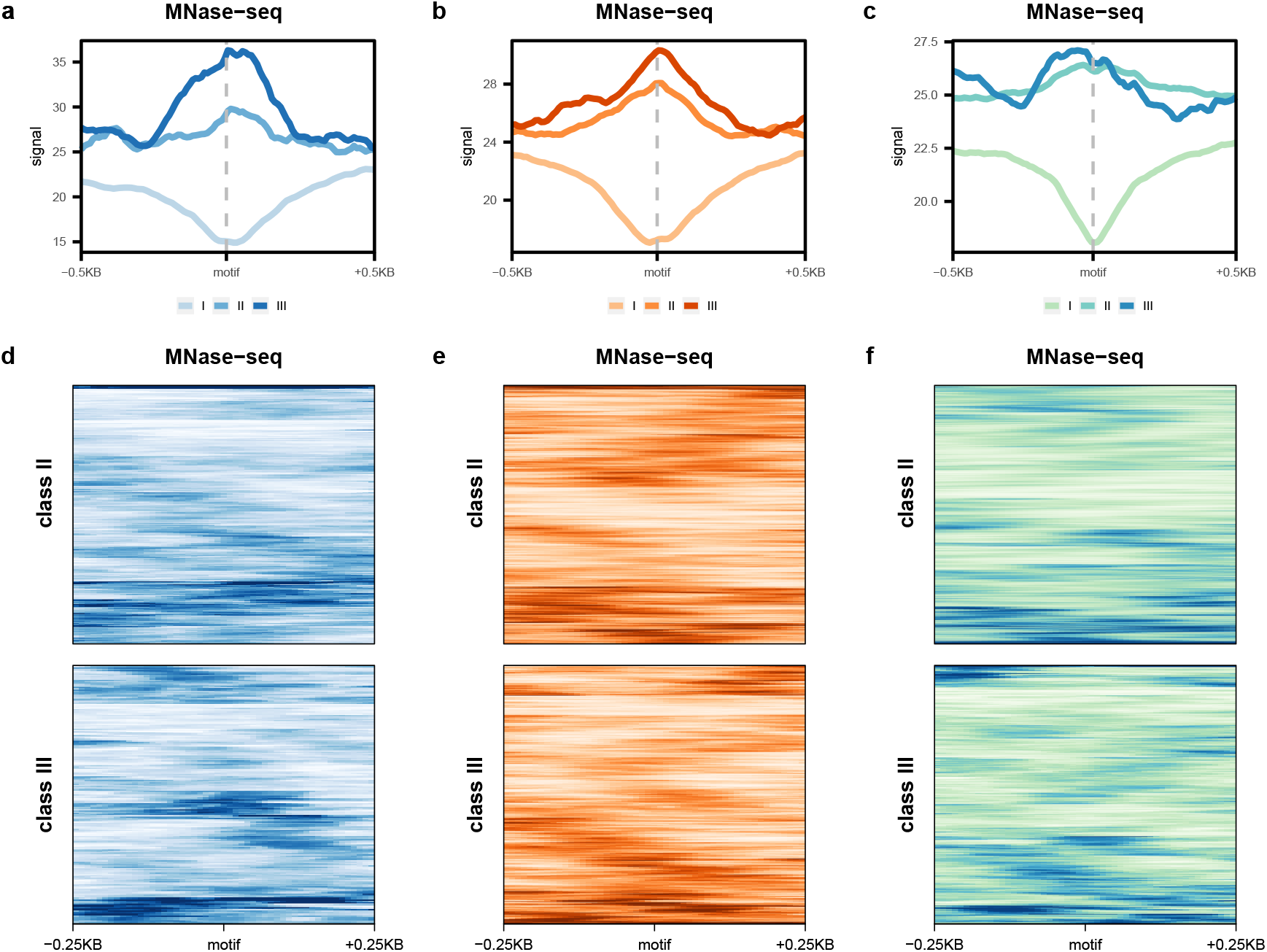
Zld, Grh and Twi do not bind preferentially to motifs in a particular position on nucleosomes. **a-c,**Metaplots showing average MNase signal from wild-type cells centered on motifs within class I, II, or III regions for Zld **(a)**, Grh **(b)**, or Twi **(c)**. **d-f,** Heatmaps showing MNase signal centered on motifs within class II and III regions for Zld **(d)**, Grh **(e)**, or Twi **(f)**. Rows are ordered based on hierarchical clustering to highlight the various patterns of MNase signal around motifs.

**Extended Data Figure 7:**
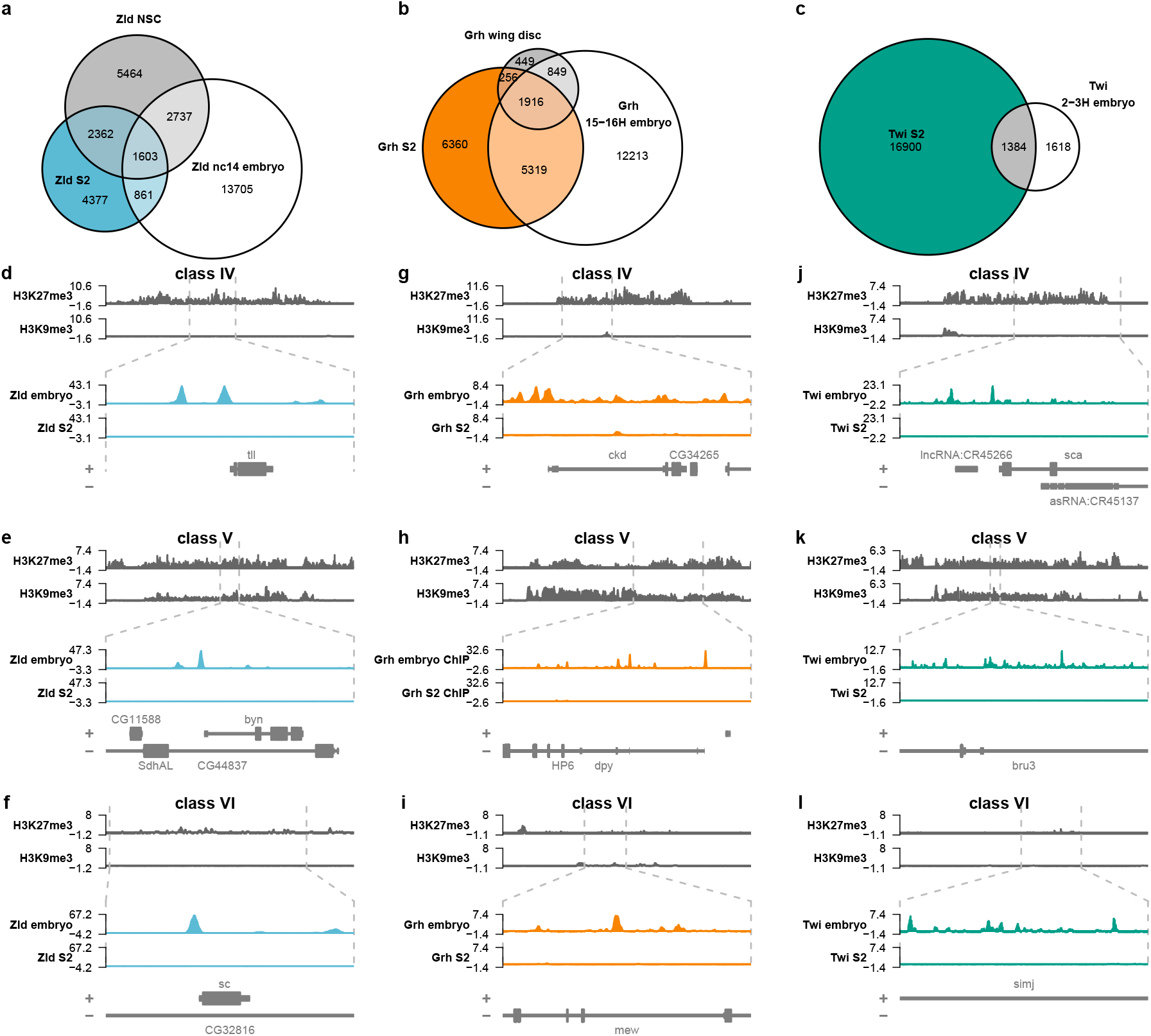
Zld, Grh and Twi display cell-type specific binding. **a-c,** Venn diagrams showing overlap between ChIP-seq peaks identified in different tissues for Zld **(a)**, Grh **(b)** or Twi **(c)**. **d-l,** Genome browser tracks showing examples of class IV, V, and VI regions for Zld **(d-f)**, Grh **(g-i)** or Twi **(j-l)**. For each example, the top tracks show H3K27me3 and H3K9me3 signal over a larger region. Dashed gray lines indicate a zoomed-in region where Zld, Grh or Twi ChIP-seq signal is shown in S2 cells or in embryos.

**Extended Data Figure 8:**
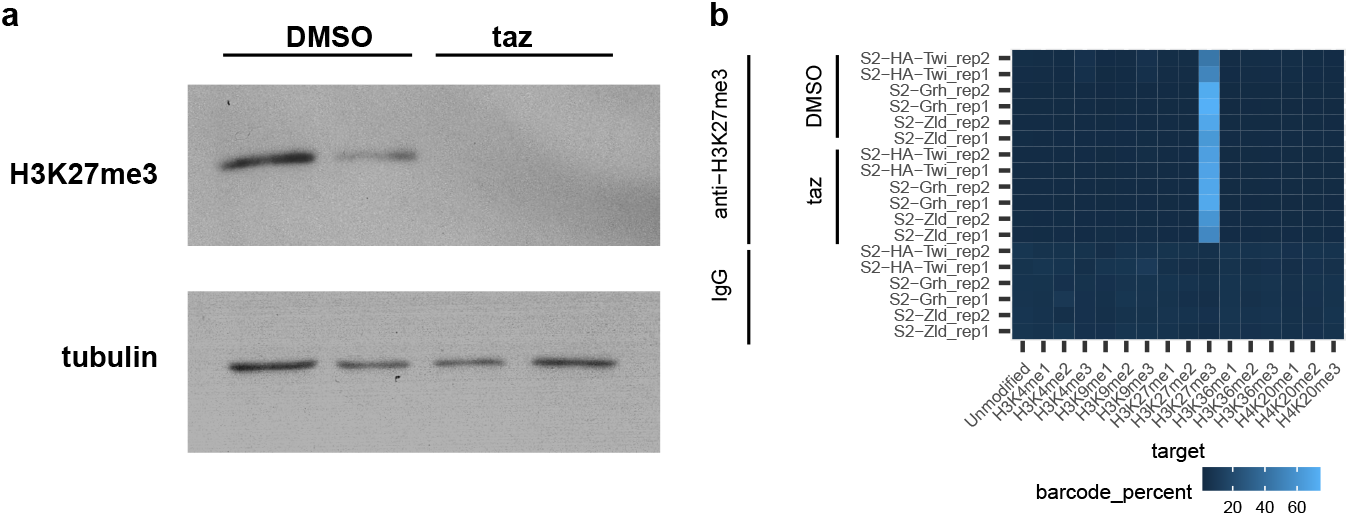
Loss of H3K27me3 in tazemetostat-treated cells. **a,** Immunoblot showing H3K27me3 levels in two replicates of DMSO- or tazemetostat-treated cells. Tubulin levels are shown as a loading control. **b,** Heatmap showing specificity of anti-H327me3 antibody in CUT&RUN reactions. A panel of barcoded spike-in nucleosomes bearing different modifications was added to each CUT&RUN reaction (see methods). For each sample, the heatmap displays the percentage of barcode reads for each sample and histone modification relative to the total number of barcode reads for all modifications.

**Extended Data Figure 9:**
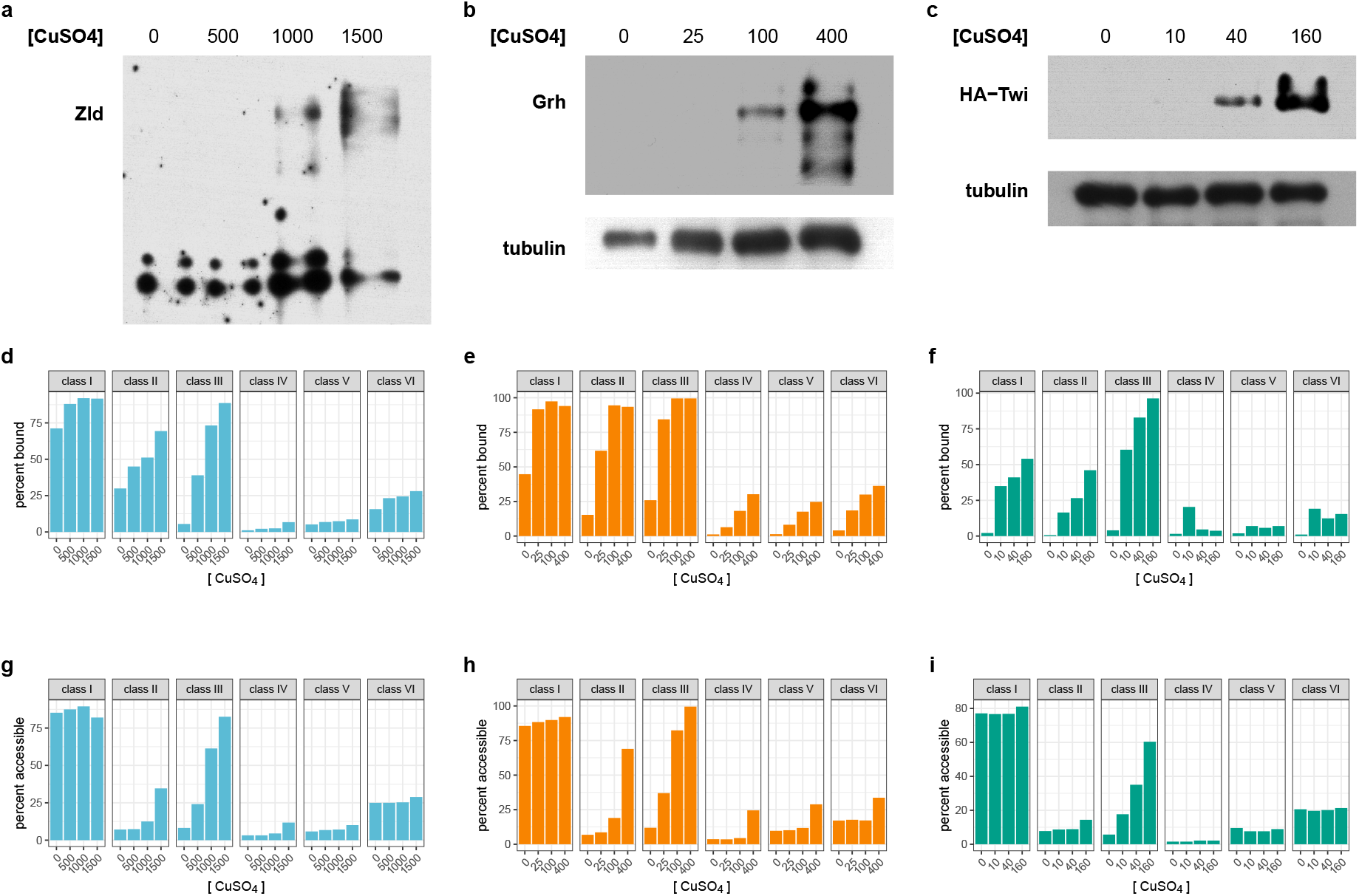
Expression of Zld, Grh, or Twi at supraphysiological levels results in chromatin opening at a small number of novel binding sites. **a-c**, Immunoblots showing Zld **(a),** Grh **(b)** or Twi **(c)** protein levels when stable cell lines are induced using different concentrations of CuSO_4_. **d-f,** Bar plots showing the percentage of previously defined class I-VI binding sites that are bound by Zld **(d)**, Grh **(e)**, or Twi **(f)** when expressed at varying concentrations. **g-i,** Bar plots showing the percentage of previously defined class I-VI binding sites that overlap an ATAC-seq peak when Zld **(g)**, Grh **(h)**, or Twi **(i)** are expressed at varying concentrations.

**Extended Data Figure 10:**
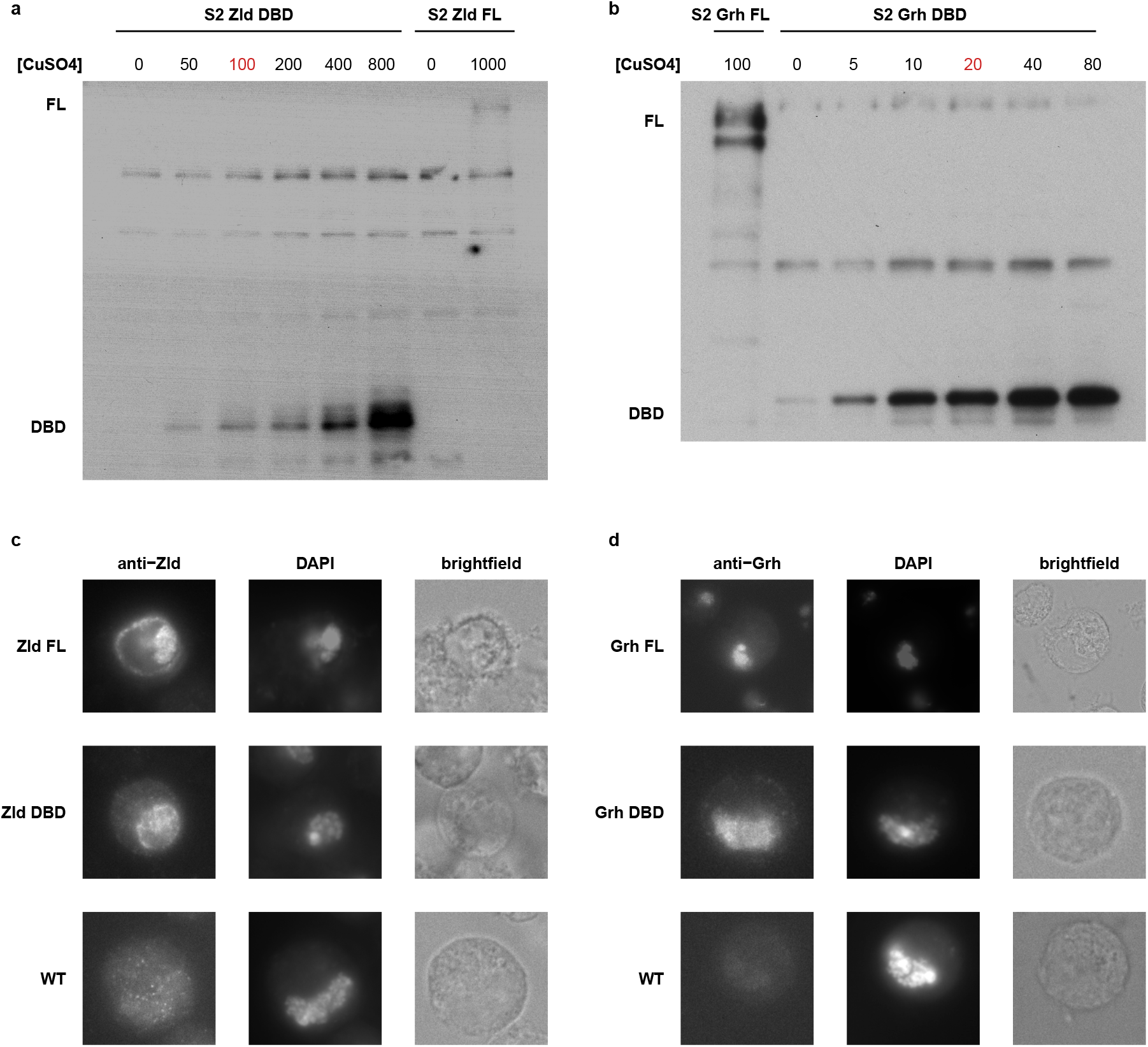
Expression of Zld and Grh DNA-binding domains at protein levels comparable to the full-length proteins. **a-b,** Immunoblots showing titration of protein levels for Zld **(a)** or Grh **(b)** DNA-binding domains to match expression of the full-length proteins. DNA-binding domain protein levels are shown at a range of CuSO_4_ concentrations and compared to an equivalent number of cells expressing full-length Zld or Grh at approximately physiological levels. The concentration of CuSO_4_ used to induce DBD expression for ChIP, ATAC and immunofluorescence experiments is indicated in red. **c-d,** Immunofluorescent microscopy images of stable cell lines expressing full-length protein or DBD only for Zld **(c)** or (Grh). Stable cell lines are compared to wild-type (WT) cells.

**Supplementary table 1: Classification of Zld, Grh and Twi ChIP-seq peaks.** Table contains information about ChIP-seq peaks for Zld, Grh and Twi, including the peak location, information about peak calling by MACS2, classification of peaks as class I, II or III, information about differential accessibility as determined by DESeq2, assignment of peaks to the nearest gene, information about differential expression of nearby genes, and motif content with each peak,

**Supplementary table 2: Previously published ChIP-seq datasets that were analyzed in this work.** For previously published ChIP-seq datasets, this table provides the target of immunoprecipitation, the cell type and/or developmental stage, the Gene Expression Omnibus accession number (GSE), and the PubMed ID for the study in which the dataset was originally published.

